# Spatial-neighbour encoding enables fast RNA 3D structure search

**DOI:** 10.64898/2026.04.19.719441

**Authors:** Ding Wang, Junru Jin, Jianbo Qiao, Leyi Wei, Shu Wu, Qiang Liu

## Abstract

Experimental and predicted RNA three-dimensional structures are expanding rapidly, but RNA structure search still lacks a compact residue-level representation that supports database-scale comparison. Using family-held-out ablations across the currently available experimental RNA structure collection, we found that spatial-neighbour features are markedly more informative for family-level discrimination than conventional backbone and base descriptors. Building on this result, we developed RiboSeek, a search framework based on a 20-letter geometric alphabet (RS-20), an 80-letter structure-and-base composite alphabet (RS-80). Across family-level classification and retrieval benchmarks, RS-80 delivered the strongest overall performance, whereas RS-20 most closely tracked US-align TM-score, indicating better preservation of geometric similarity. RiboSeek searches the full experimental RNA structure database in 204 ms per query and can be applied to predicted RNA structure libraries to prioritize candidate structural relationships for downstream analysis.

## Introduction

Experimental and predicted RNA three-dimensional structures are accumulating rapidly. As these resources grow, exhaustive pairwise structural alignment becomes too slow for routine database-scale search, creating a need for compact representations that support fast and sensitive comparison. In proteins, Foldseek showed that encoding each residue as a discrete structural token can convert structure comparison into sequence-style alignment and enable interactive search across large structure collections[3, 4]. An equivalent infrastructure has not yet been established for RNA.

A fifteen-year lineage of RNA structural alphabets — SARSA[5], iPARTS[6], R3D-BLAST[8], iPARTS2[7] and R3D-BLAST2[9] — has largely encoded per-residue geometry through backbone pseudo-torsions. Despite differences in implementation, none of these alphabets explicitly represents the geometry of non-sequential spatial neighbours around each residue. This omission is notable because many tertiary features of RNA structure are defined less by local backbone conformation than by the spatial arrangement of nearby bases and residues. Whether an explicitly neighbour-aware design can improve compact RNA structure encoding has remained unclear.

We therefore asked two related questions. First, how much information do spatial neighbours contribute beyond backbone descriptors for RNA structure search? Second, can that information be compressed into a compact alphabet that supports rapid database-scale search without sacrificing structural fidelity? To address these questions, we compared alphabets trained on nested feature sets under a family-held-out protocol and incorporated the resulting representations into a C-accelerated search pipeline.

Here we present RiboSeek, an RNA structure-search framework built around two complementary alphabets: RS-20, a 20-letter geometric alphabet, and RS-80, an 80-letter composite that combines structure with nucleotide identity. We show that spatial-neighbour features dominate performance in family-held-out ablations, that the resulting framework improves family-level classification and retrieval relative to existing RNA structure-search baselines, and that RS-20 more closely tracks US-align TM-score. We further illustrate how the framework can be applied to predicted RNA structure libraries for hypothesis-generating analyses.

## Results

We first used family-held-out ablations to determine which residue-level features are most informative for compact RNA structure encoding and to define the production alphabets RS-20 and RS-80. We then evaluated the resulting framework on family classification, retrieval and geometric-fidelity benchmarks before applying it to predicted RNA structure libraries. An overview of the system is given in Figure 1.

**Figure 1:**
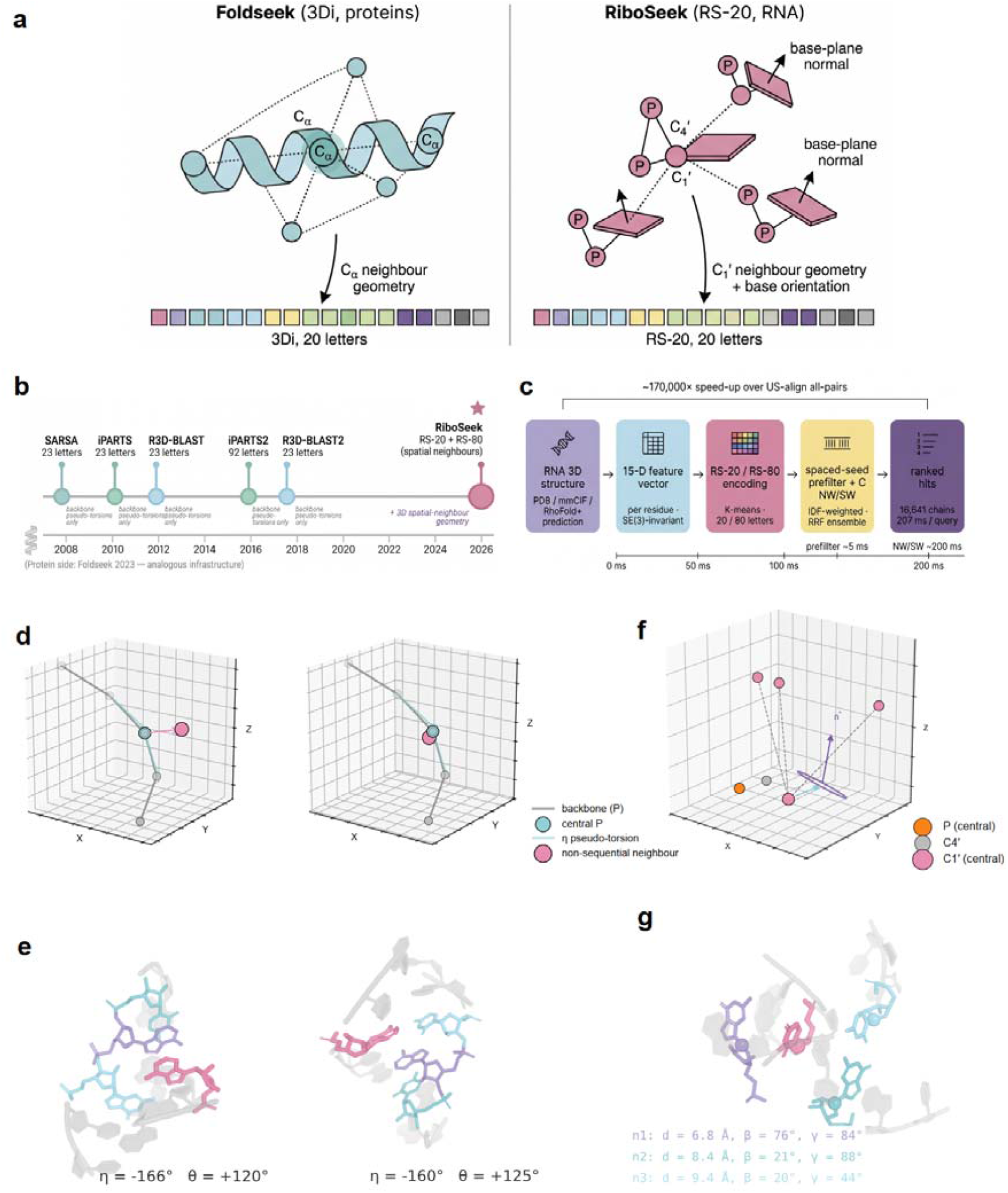
Overview of RiboSeek. (a) Schematic contrast between the Foldseek 3Di alphabet for proteins (Cα neighbours → 20 tokens) and the RiboSeek RS-20 alphabet for RNA (C1’ neighbours + base-plane orientation → 20 tokens). (b) Published RNA structural alphabets (2008–2017) encoded per-residue geometry through backbone pseudo-torsions; RS-20 explicitly encodes the geometry of non-sequential spatial neighbours. (c) End-to-end pipeline: RNA 3D structure → 15-D feature vector → RS-20 / RS-80 encoding → spaced-seed prefilter + C-accelerated NW/SW → ranked hits. (d, e) A schematic illustration (d) and two real PDB residues whose backbone η/θ pseudo-torsions match to within 7.6° but whose top-3 spatial-neighbour arrangements differ substantially (e). (f, g) The spatial-neighbour descriptors used by RS-20, shown schematically (f) and for a real residue (g): top-3 non-sequential C1’ neighbours × {distance, base-normal angle, C1’-centroid-direction angle} + base-plane orientation + sequential inter-residue distances + non-sequential contact count. Panels (e) and (g) are PyMOL renderings from the mini-PDB fragments used during alphabet construction (Methods § Feature extraction).

### Spatial-neighbour features carry more same-family signal than backbone-centric features

To test which residue-level descriptors are most useful for RNA structure search, we trained K-means alphabets (K = 20) on seven nested feature sets extracted from the 15,391 production RNA chains and evaluated them under a family-held-out protocol. In this protocol, K-means is trained only on chains whose Rfam annotations do not appear in the 11,170-pair hard-negative benchmark (3,419 training chains), minimizing family-level information leakage into codebook construction.

The resulting ablation (Fig. 2a; full numerical table in Supplementary Table S10) showed a clear ordering of feature utility. A three-feature set encoding spatial-neighbour geometry alone (AUC = 0.945; 95 % bootstrap interval 0.940–0.950) outperformed the combined seven-feature backbone-and-base set (AUC = 0.921; 0.915–0.926); the two intervals do not overlap. Adding spatial-neighbour features on top of the traditional set gave no further gain, and removing backbone η / θ from the combined 15-D representation gave a small additional improvement (AUC 0.955). Together, these comparisons indicate that the main design gain comes from encoding spatial-neighbour geometry, whereas backbone pseudo-torsions contribute little once neighbour features are included. Accordingly, the production representation used below centres on spatial-neighbour geometry together with base-related and sequential descriptors, rather than on the pseudo-torsions that defined five prior published RNA alphabets (Supplementary Table S1).

**Figure 2:**
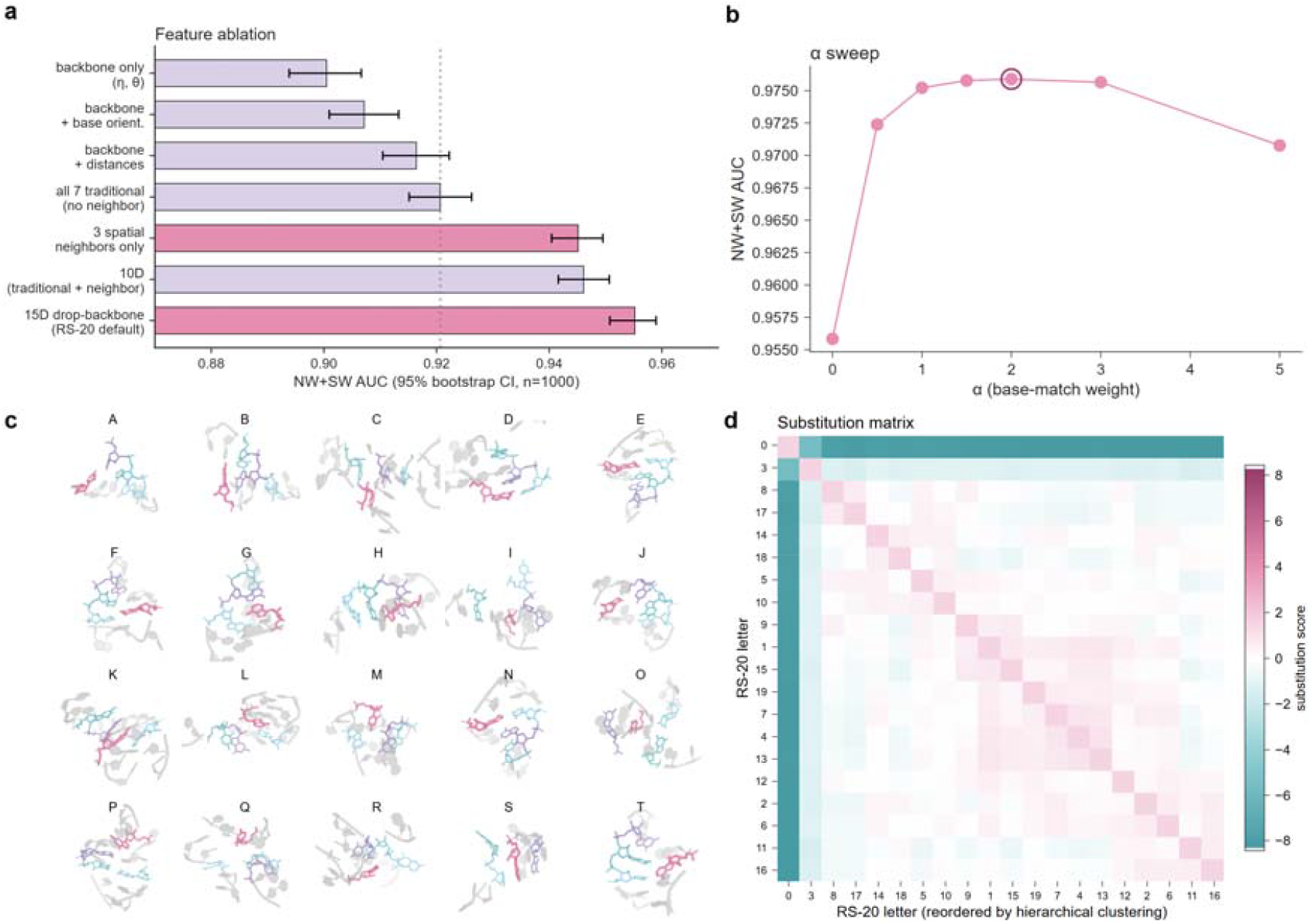
Feature ablation and RS-20 alphabet. (a) Feature-ablation bar plot with 95 % bootstrap CIs (n = 1,000) on NW+SW AUC. Three spatial-neighbour features alone reach AUC = 0.945 [0.940, 0.950], exceeding all seven traditional backbone/base features combined (0.921 [0.915, 0.926]); intervals do not overlap. (b) α-weight sweep for the RS-80 composite alphabet (α = 2 optimal), balancing the structural and sequence contributions. (c) Representative 3D microenvironments for all twenty RS-20 clusters (labelled A–T; rose = centre nucleotide; lavender, mint, sky = top-3 spatial neighbours; grey = local sequence context). (d) RS-20 substitution-matrix heatmap, hierarchical-cluster-reordered, illustrating the structure that the K-means alphabet learns over geometric features. Alphabet-size K □ {10, 15, 20, 25, 30, 40} and top-k-neighbour k □ {1, …, 5} hyperparameter sweeps (K = 20 plateau at K ≥ 15; top-3 saturation) are shown in Supplementary Fig. S1.

### Two complementary alphabets for geometric fidelity and classification

On the basis of the ablation, we defined two complementary alphabets. **RS-20** is a 20-letter K-means alphabet over the 15-D production feature vector and is designed to preserve structural similarity. **RS-80** augments the same structural states with nucleotide identity to improve family-level discrimination (Methods § RS-20 and RS-80 encodings). The two encoders therefore serve distinct but related purposes: RS-20 prioritizes geometric fidelity, whereas RS-80 prioritizes retrieval sensitivity.

Parameter sweeps supported the chosen design. Performance plateaued at alphabet size K = 20 and saturated when the top three non-sequential neighbours were included (Supplementary Fig. S1). For RS-80, α = 2 provided the best balance between structural and nucleotide-identity contributions on the hard-negative benchmark (Fig. 2b). We therefore use RS-20 for geometry-oriented analyses and RS-80 for family-level search and retrieval.

### RiboSeek outperforms existing RNA structure-search methods

We next compared RiboSeek with published RNA structure-search methods and sequence-based baselines on the 11,170-pair length-matched hard-negative Rfam benchmark. RS-80 attained the highest overall AUC (0.935), followed by RS-20 (0.898); both exceeded BLAST (0.860), iPARTS2 (0.860), R3D-BLAST2 (0.835) and the Foldseek pseudo-protein control (0.873) (Fig. 3a, left). RS-20 achieves its result with only 20 structural states, indicating that neighbour-aware encoding improves family-level discrimination under a benchmark designed to reduce simple length-based shortcuts. Stratification by Needleman–Wunsch pairwise identity, which is more robust to indels than k-mer Jaccard (Supplementary Methods § S1), localises the advantage of structural search (Fig. 3a, right). At 50–70 % identity (n = 171 pairs) BLAST AUC decreased to 0.60 whereas RS-80 remained at 0.95; above 70 % identity both methods approach their ceiling values.

**Figure 3:**
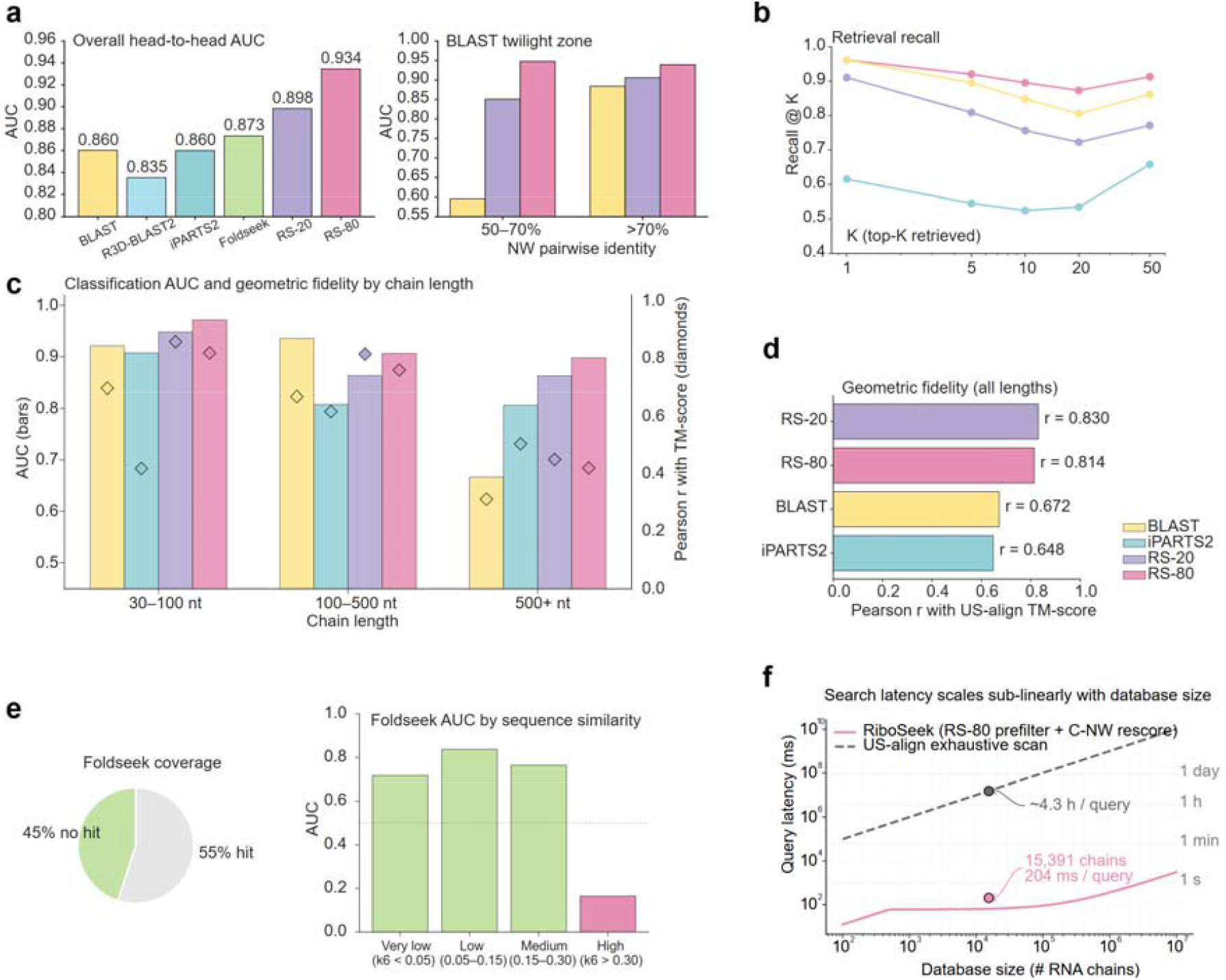
Head-to-head benchmark, retrieval and geometric fidelity. All panels use 11,170 length-matched hard-negative Rfam pairs (8,170 same-family positives + 3,000 length-matched different-family negatives) unless otherwise stated. (a) Method head-to-head. *Left*: overall AUC across six methods — BLAST 0.860, R3D-BLAST2 0.835, iPARTS2 0.860, Foldseek pseudo-protein control 0.873, RS-20 0.898, RS-80 0.935. *Right*: NW-pairwise-identity stratification (BLAST twilight zone). (b) Retrieval recall-at-K curves on 156 Rfam-representative queries against a 1,041-chain retrieval database. (c) Classification AUC (bars, left axis) and Pearson r with US-align TM-score (diamonds, right axis) stratified by chain length; RS-20 attains the highest geometric fidelity across all strata. (d) Geometric fidelity: overall Pearson r with US-align TM-score on 600 chain pairs — RS-20 0.830, RS-80 0.814, BLAST 0.672, iPARTS2 0.648. (e) Failure modes of the Foldseek pseudo-protein control: 45 % of pairs receive no hit, and within the high-sequence-similarity stratum the AUC is 0.165. A four-mapping sensitivity control (Supplementary Fig. S4) indicates that this behaviour is determined by the 3Di encoder’s protein-geometry training distribution. (f) Search latency as a function of database size; 204 ms per query on the 15,391-chain production database.

In the retrieval setting (156 queries × 1,041 target chains, two queries per Rfam family), RS-80 achieved recall@1 = 0.962, recall@10 = 0.896 and recall@50 = 0.913 (Fig. 3b), matching or exceeding the sequence-based BLAST baseline across all K. RS-20 remained competitive (recall@10 = 0.757) despite encoding no sequence information. Length-stratified analyses showed that the gap over sequence-based search widens for longer RNAs, and RS-20 attains the highest Pearson correlation with US-align TM-score in the 30–100 nt range (r = 0.857; Fig. 3c). On a held-out set of 600 chain pairs, RS-20 attained the highest overall Pearson correlation with US-align TM-score (Fig. 3d; RS-20 0.830, RS-80 0.814, BLAST 0.672, iPARTS2 0.648), consistent with its role as a geometry-focused representation. These results indicate that RiboSeek is particularly useful in the regime where sequence similarity alone becomes difficult to exploit.

An RNA-adapted Foldseek control did not transfer well in this setting (Fig. 3e): 45 % of benchmark pairs received no hit, and the high-sequence-similarity stratum showed AUC = 0.165 (anti-correlated). The same failure pattern was observed across four orthogonal residue-mapping controls (mnemonic, size-matched, polarity-matched and random; Supplementary Fig. S4 and Table S4), indicating that the poor transfer arises from applying a protein-geometry-trained encoder to RNA local geometry rather than from a particular residue-mapping choice. At runtime, exhaustive C-accelerated NW search of the 15,391-chain production database took 22.3 s per query on a single thread, whereas the spaced-seed prefilter reduced this to 204 ms per query — an approximately 75,000-fold speed-up relative to exhaustive US-align of the same database (Fig. 3f).

### RS-20 recovers cross-family similarities within Rfam clans

To examine whether the geometric alphabet captures biologically meaningful similarities beyond Rfam-level labels, we compiled 387 cross-family within-Rfam-clan chain pairs with US-align TM ≥ 0.40 and BLAST *e*-value > 0.01 — pairs that are structurally similar but not recovered by sequence search — spanning six major Rfam clans (RNase P, SRP RNA, tRNA / tmRNA, LSU rRNA, purine riboswitches and SAM riboswitches; Fig. 4a, Supplementary Fig. S8, Supplementary Table S8).

**Figure 4:**
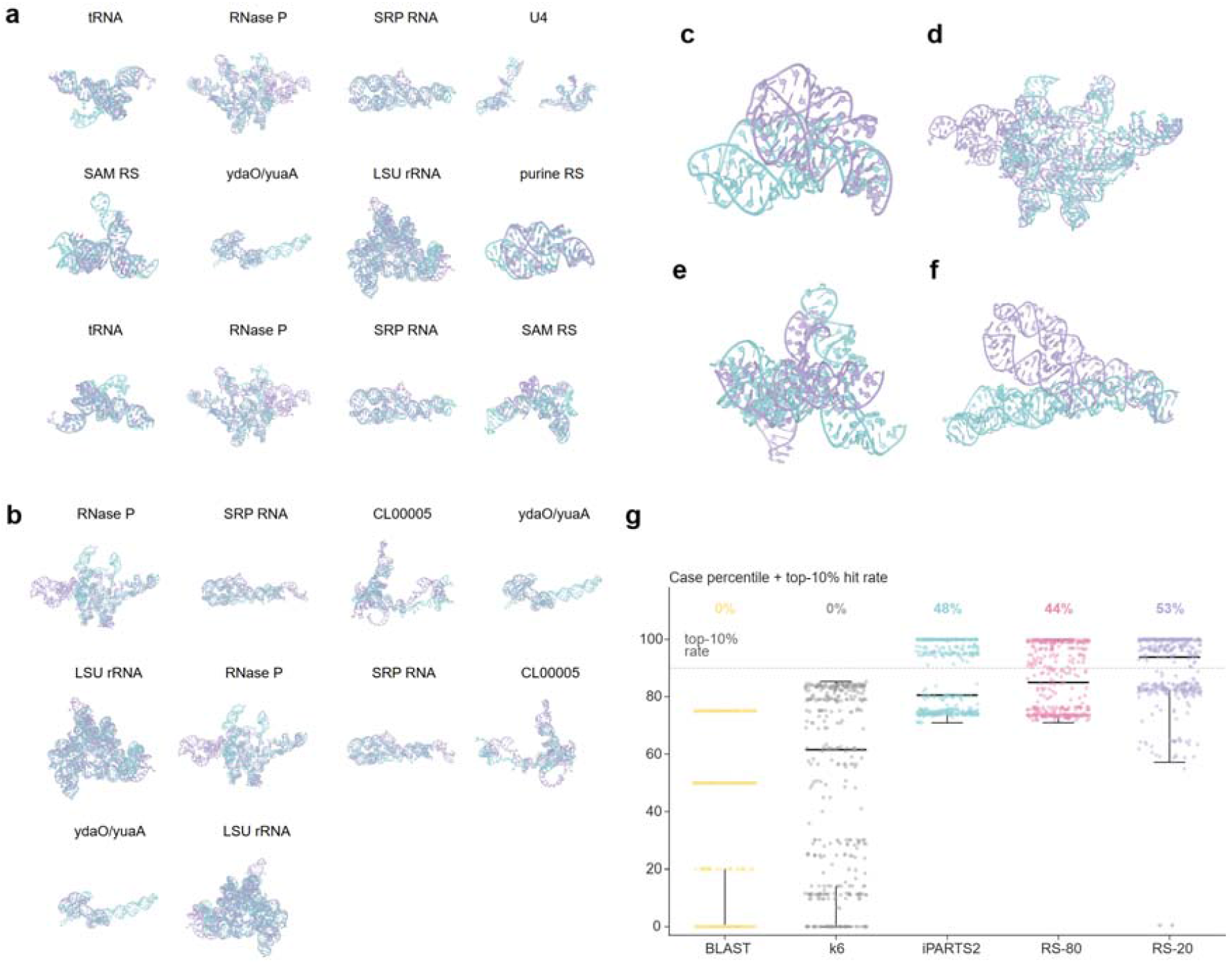
Cross-family within-clan case studies. (a) Twelve cross-family within-Rfam-clan case superpositions (coverage view across six clans: RNase P, SRP RNA, tRNA / tmRNA, LSU rRNA, purine riboswitches and SAM riboswitches). Each panel shows the C1’ backbone trace of two chains after US-align rotation; full metadata per panel is given in Supplementary Fig. S8. (b) Twelve within-clan case pairs for which RS-20 (20 letters) places the pair at a higher percentile of a cross-family noise distribution than iPARTS2 (92 letters); the largest gap is +14 percentile points. Full annotation is in Supplementary Fig. S9. (c–f) Four cases rendered as PyMOL cartoons after US-align rotation, spanning the purine riboswitch (CL00123), RNase P (CL00002), SAM riboswitch (CL00012) and SRP RNA (CL00003) clans. (g) Percentile-rank distribution of all 387 BLAST-fail TM ≥ 0.40 cases across five methods; the fraction reaching the 90th percentile of a 3,000-pair cross-family noise distribution is labelled above each strip.

Representative cases span several well-studied RNA classes: an adenine- and guanine-bound aptamer sharing a three-way-junction fold (purine riboswitch; Fig. 4c), bacterial type-A and type-B RNase P RNAs sharing the catalytic core (Fig. 4d), members of different SAM riboswitch classes sharing a SAM-binding pocket geometry (Fig. 4e), and the Alu domain of signal-recognition-particle RNA in archaea and bacteria (Fig. 4f). In these cases, RS-20 preferentially ranked the shared structural cores despite sequence divergence or differences in peripheral elements. Across all 387 cases (Fig. 4g), the BLAST and k6 Jaccard baselines do not distinguish individual pairs from cross-family noise (0 % reach the 90th percentile of the noise distribution, by construction since pairs were selected as BLAST-fail). iPARTS2, RS-80 and RS-20 reach the top 10 % of noise in 48 %, 44 % and 53 % of cases respectively. Together, these within-clan examples provide qualitative support for the benchmark results by showing that the neighbour-aware encoding retrieves structurally interpretable relationships in known RNA systems.

### Dark-family search nominates candidate structural relationships

We next asked whether RiboSeek could be used to search predicted RNA structure libraries without experimental representatives. Using RhoFold+[2] in single-sequence mode, we generated predictions for 3,694 dark Rfam families (lengths 40–200 nt, AUGC only), retaining 3,680 that completed prediction and encoding. On 419 predictions for which an experimental reference was available, the median predicted-vs-experimental US-align TM-score is 0.40 (Fig. 5b), consistent with the reported performance of current single-sequence RNA structure predictors[2] but below the 0.50 “same-fold” threshold. Within this limitation, retrieval in the operational predicted-query → predicted-database setting was substantially more reliable than direct predicted-query → experimental-database retrieval (RS-80 recall@10 = 0.90 versus RS-20 recall@10 = 0.22; Fig. 5c), consistent with the alphabet producing self-consistent assignments under a given predictor’s systematic bias.

**Figure 5:**
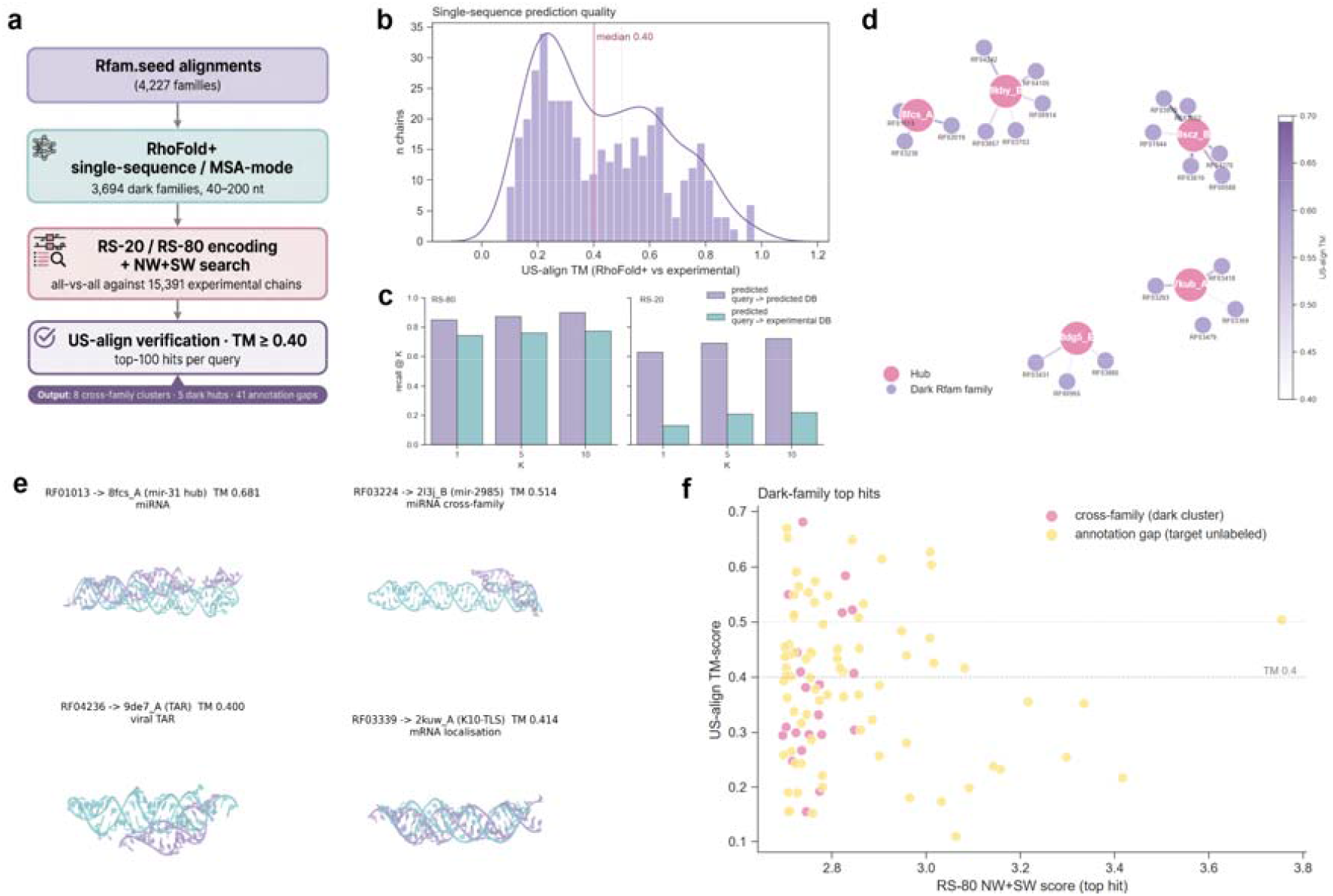
Dark-family search and recurrent hub-like matches. (a) Predicted-library pipeline: Rfam.seed → RhoFold+ → RS-20 / RS-80 encoding → NW+SW search → US-align verification. (b) RhoFold+ single-sequence prediction quality on 419 Rfam chains with an experimental reference (median US-align TM = 0.40). (c) Predicted-library retrieval: predicted-query → predicted-database (operational setting) compared with predicted-query → experimental-database (cross-domain diagnostic). (d) Network of recurrent hub-like matches: five experimental chains each receive three or more different dark Rfam families as the closest structural hit; node and edge colour encode US-align TM. (e) Four candidate cross-family superpositions after US-align rotation (the complete set of eight is listed in Supplementary Table S6). (f) RS-80 score against US-align TM for the top-100 dark-family hits; the eight candidate cross-family pairs and 41 annotation-gap hits to unlabelled experimental chains are highlighted.

Among the top-100 verified hits, US-align supported 49 chain pairs with TM ≥ 0.40 and 23 with TM ≥ 0.50. Only **8** of these 49 were strict cross-family cases, in which the top-ranked experimental chain belonged to a different Rfam family from the dark query. Most of these candidate pairs involved short hairpin-like RNAs spanning several functional categories that are kept separate by sequence-based family definitions (full list in Supplementary Table S6). Re-predicting the eight dark queries in MSA mode increased the median TM-to-match from 0.48 to 0.53 and the TM ≥ 0.5 rate from 3/8 to 5/8 (Supplementary Methods § S9), indicating that the similarity is not solely attributable to single-sequence prediction noise. Given the small number of cases and the reliance on predicted structures, these should be considered candidate cross-family structural relationships rather than established family reassignments.

A stronger signal emerged from recurrent hub-like matches, in which the same experimental chain appeared as the closest structural hit for multiple dark families. Using a threshold of at least three converging dark families, we identified five such hubs (Fig. 5d). Across 48 additional seeds sampled from the 19 implicated dark families, 46 of 47 evaluable seeds (98 %) retained TM ≥ 0.40 to the expected hub and 33 of 47 (70 %) retained TM ≥ 0.50 (Supplementary Fig. S11). In a matched negative control, the same pipeline applied to four non-dark Rfam families with distinct known folds (three seeds each; 12 total) produced 12 distinct experimental top hits (Supplementary Methods § S6), arguing against a trivial predictor-specific fold-attractor explanation. The biological interpretation of individual hubs will nonetheless require independent validation.

In addition to the candidate cross-family pairs, the search highlighted 41 TM ≥ 0.40 hits to experimental chains that are unlabelled in the current Rfam–PDB mapping, eight of which involve a single recurrent chain (8scz_B). Four of the five dark hubs remain unannotated under the current Rfam 15.x PDB mapping. These results are best treated as candidate annotations and hypotheses for follow-up curation.

### A global view of the RS-20 representation space

To place the experimental and predicted chains in a shared representation space, we embedded their RS-20 1+2-gram profiles into two dimensions with UMAP. Major Rfam clans occupied largely separated regions, and the five recurrent dark hubs from Fig. 5d localised to a common area of the map (Fig. 6). This embedding provides a global visual summary of the similarity structure already seen in the pairwise search results.

**Figure 6:**
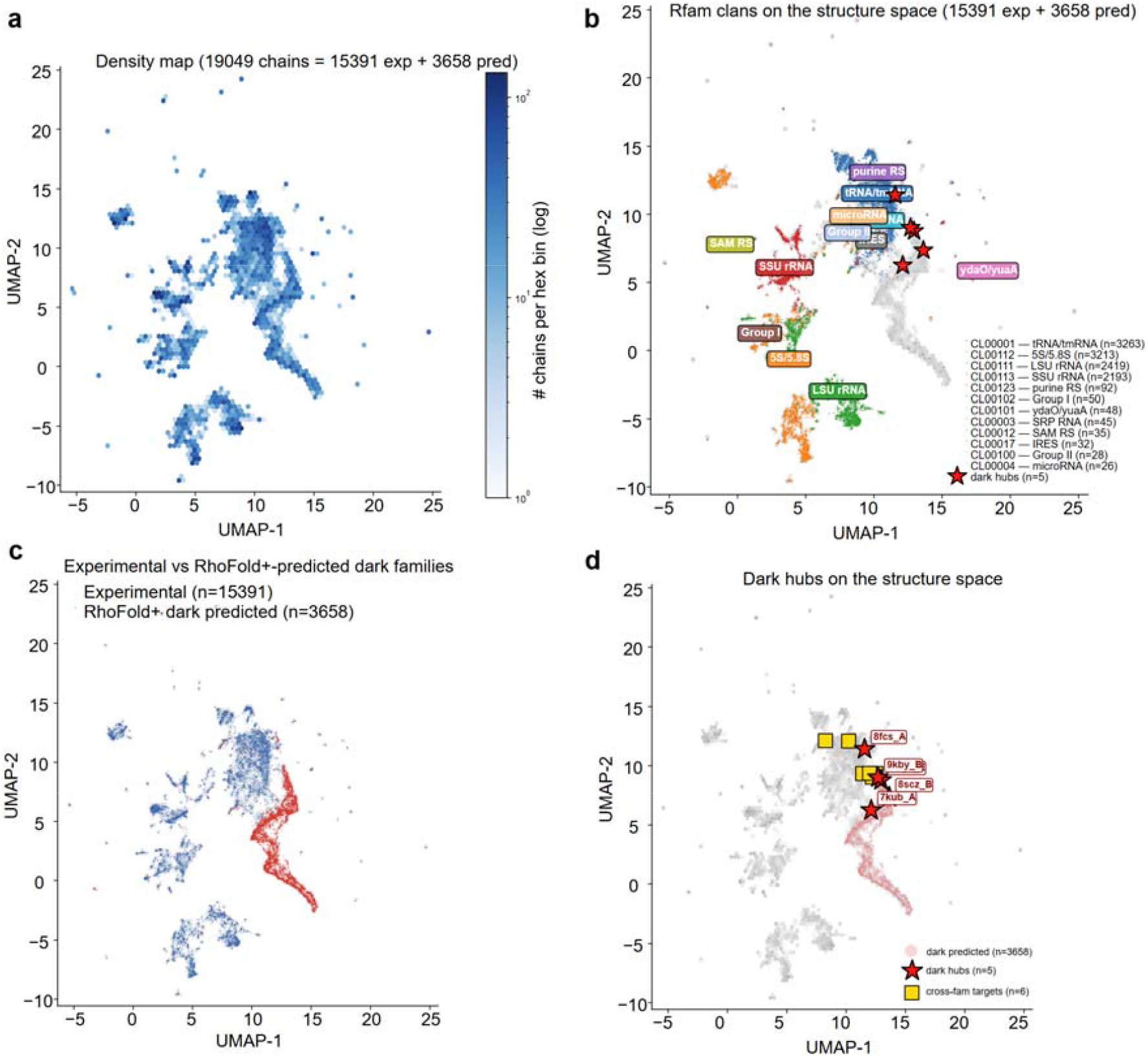
Global RS-20 representation space. UMAP embedding of 19,049 RNA chains (15,391 experimental + 3,680 RhoFold+-predicted dark-family), each represented by its 420-dimensional RS-20 1+2-gram profile (cosine metric, n_neighbors = 15, min_dist = 0.1). (a) Hexbin density. (b) Coloured by Rfam clan; major clans (CL00001 tRNA/tmRNA, CL00111 LSU rRNA, CL00112 5S rRNA, CL00123 purine riboswitches, CL00012 SAM riboswitches, CL00003 SRP RNA) occupy largely separated regions. (c) Experimental chains versus RhoFold+ predictions. (d) The five dark hubs from Fig. 5d localise to a common region of the embedding.

## Discussion

The central result of this study is that compact RNA structure search benefits more from encoding non-sequential spatial-neighbour geometry than from further refining backbone-centred residue descriptors. This finding motivated two complementary encoders — RS-20 for geometric fidelity and RS-80 for family-level discrimination and retrieval — and a search framework that performs strongly across classification, retrieval and runtime benchmarks.

Earlier RNA structural alphabets describe each residue primarily through local backbone conformation. Our ablation indicates that this is not the most informative choice once non-sequential spatial context is encoded explicitly. In RNA, many tertiary interactions — including long-range base stacking, A-minor motifs and kissing-loop contacts — are defined by the relative arrangement of nearby residues and bases rather than by backbone geometry alone. The stronger performance of neighbour-aware encodings therefore supports a shift from backbone-centred to context-centred residue representations. Foldseek reported an analogous observation for proteins using a learned VQ-VAE over Cα-neighbour geometry; the same principle appears to operate in RNA under a simpler hand-designed representation. The two alphabets serve distinct roles: RS-20 is the more faithful geometric representation and attains the highest Pearson correlation with US-align TM-score in our benchmarks, whereas RS-80 adds nucleotide identity as a separable channel and is correspondingly better suited to family-level discrimination and retrieval. A compact RNA structural alphabet may also provide a useful starting point for future structure-aware RNA representation learning[31, 33–36], although that application lies beyond the scope of the present study.

In its current form, RiboSeek is most useful for small-to medium-sized RNAs in the 30–500 nt range emphasized by our benchmarks, including tRNAs, miRNA precursors, riboswitches, ribozymes, and many SRP- and RNase P-class functional RNAs. In this regime, the framework provides strong retrieval performance and good agreement with structure-based similarity measures. Performance declines on substantially longer and more clearly multi-domain RNAs, for which a single global or local alignment is unlikely to capture the relevant structural organization. The current predicted-library application is likewise constrained by the quality of available RNA structure predictors, especially in single-sequence mode, and the production spaced-seed prefilter still trades maximal recall for low latency. The predicted-library analysis also raises the possibility that sequence-defined RNA families can, in some cases, partition structure space more finely than fold-level similarity would suggest. Our data are consistent with this possibility for a small number of short hairpin-like systems, but the evidence remains provisional because it depends on predicted structures and yields only a limited number of strict cross-family cases. In this context, the main value of RiboSeek is to generate candidate structural relationships for follow-up study. These limitations define the present scope of the method but do not alter the main benchmark observations reported here.

## Methods

### Data

*RNA structures come from the PDB and are filtered for complete per-residue features. Several distinct subsets are used for training, benchmarking, retrieval and predicted-library search, and are defined at first mention below*.

RNA structures were obtained from the RCSB Protein Data Bank (April 2026, 9,715 RNA-containing entries; 2,241 RNA-only, 7,474 RNA–protein complexes). From 9,701 downloaded mmCIF files we parsed 25,218 RNA chains, of which **15**,**391** yielded complete 15-D feature vectors (§ Feature extraction) across ≥ 50 % of residues; this set is used throughout as the production experimental database (also referred to below as the search database). Rfam family membership was taken from Rfam 14.10 (`Rfam.pdb` flat file), yielding 102 families with ≥ 3 chains in our dataset; the **benchmark sub-pool of 6**,**433 chains** is the subset of the 15,391 that belongs to one of those 102 benchmarked families. Throughout the Results, three further subsets are used and defined at first mention: the **11**,**170 length-matched hard-negative pair** classification benchmark (§ Benchmarks); a **1**,**041-chain retrieval database** with at most 20 chains per family (§ Benchmarks); and a **3**,**419-chain family-disjoint training pool** used for codebook fitting (§ Family-aware K-means).

### Feature extraction

*Each nucleotide is represented by invariant descriptors of local backbone geometry, sequential inter-residue distances, base orientation and non-sequential spatial neighbourhood. All descriptors are constructed from distances or angles and are therefore invariant to rigid-body rotation and translation*.

For each nucleotide *i* in a chain of length *L* ≥ 5, we extract a 17-dimensional feature vector **x**_*i* from the atomic coordinates. The features partition into four groups:

#### Backbone pseudo-torsions (2-D)

Following Wadley et al.[20]:

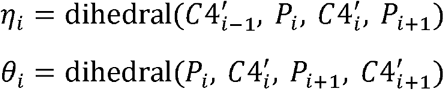

#### Sequential inter-residue distances (3-D)

*d*_pp_ = || *P*_*i*+1_ − *P*_*i*_||, 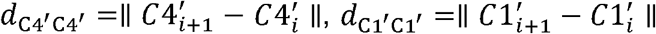.

#### Base-plane descriptors (2-D)

The base-plane normal 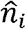 is the smallest-eigenvalue eigenvector of the SVD of the base-heavy-atom coordinate covariance. The stacking angle is 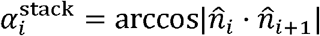; the glycosidic angle is 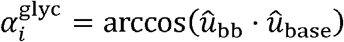 where 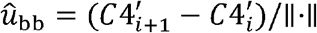 and 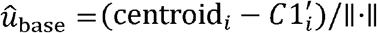.

#### Spatial-neighbour descriptors (9-D = 3 features × top-3 neighbours)

For each of the three nearest non-sequential C1’ atoms (|*i* □ *j*| > 2) within 12 Å, identified by a *k*-d tree query, we record: (a) distance 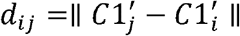, (b) base-normal angle 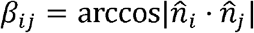, (c) base-centroid direction angle 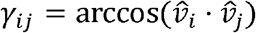 where 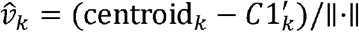.

#### Contact count (1-D)

the number of C1’ atoms within 10 Å of 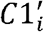, excluding *i ±* 2.

All 17 features are rotation- and translation-invariant scalars (distances or angles computed from dot products; no cross products), making the encoding SE(3)-invariant by construction (Supplementary Methods S10).

The 15-D production feature vector used for RS-20 drops η and θ (dimensions 0– 1), retaining dimensions 2–16. This choice was the empirical winner of the ablation (Fig. 2a): adding backbone pseudo-torsions to the spatial-neighbour features degrades AUC.

### Family-held-out K-means

*To avoid family-level information leakage, the codebook is trained only on chains outside the benchmark families*. K-means training excludes every chain annotated to any Rfam family that participates in the benchmark, leaving 3,419 of the 15,391 production chains as a training pool. From this pool we subsample up to 2,000 chains, concatenate their per-residue feature vectors, and normalise each dimension to zero mean and unit variance (statistics computed on training data only). We then fit scikit-learn `MiniBatchKMeans` with K = 20 codebook entries, batch size 10,000, 200 iterations, and random seed 42.

The substitution matrix ***M*** is derived from inter-centroid Euclidean distances:

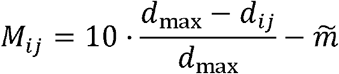

where *d*_*ij*_ =|| ***c***_*i*_ − ***c***_*j*_ ||_2_, *d*_max_ = _*i≠j*_ *d*_*ij*_, and 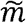 is the median of the off-diagonal entries, serving as a zero-point re-centring that ensures unrelated letter pairs receive negative scores on average.

### RS-20 and RS-80 encodings

*RS-20 assigns each residue to its nearest structural centroid. RS-80 augments the same structural states with nucleotide identity, allowing geometry and sequence to contribute separately but jointly to alignment scoring*.

RS-20 encodes each residue as its nearest K-means centroid index (1 of 20). RS-80 concatenates with nucleotide identity: J(i) = RS(i) × 4 + base(i) □ {0, …, 79}, with base(i) □ {A = 0, C = 1, G = 2, U = 3}. The RS-80 substitution matrix is M_J[i, j] = M_RS[□i/4□, □j/4□] + α · M_base[i mod 4, j mod 4], with M_base a diagonal-heavy 4 × 4 matrix (+2 on diagonal, −1 off). The weight α = 2 was selected by a single-parameter sweep on the hard-negative benchmark (Fig. 2b).

### C-accelerated alignment

*Alignment is implemented in C for speed and combined as a normalized global + local score per query–target pair*.

Global (Needleman–Wunsch[14]) and local (Smith–Waterman[15]) alignment are implemented in C and exposed to Python through `ctypes`. The NW recurrence is:

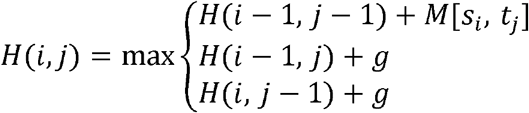

with *g* = −2 (linear gap penalty) and boundary conditions *H*(*i*, 0) = *ig, H*(0, *j*) = *jg*. The SW variant adds a fourth branch *H*(*i, j*) ≥ 0. Only two DP rows are retained (*O*(*m*)memory). Compilation uses `-O3 -march=native`; a single 150-nt × 150-nt NW alignment takes ∼120 *µ*s on one CPU core (∼354× over an equivalent Python loop).

Raw NW and SW scores are each z-score normalised across all database chains for a given query:

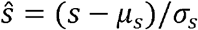

and summed to yield the composite NW+SW score used throughout.

### Spaced-seed prefilter

*The prefilter is a fast candidate-generation stage: it narrows the set of database chains that are passed to exhaustive NW / SW rescoring. Its recall is therefore reported separately from the end-to-end retrieval recall in the main text, which refers to the final ranking produced after rescoring*.

For database search we precompute inverted indices over three spaced-seed patterns on the RS-80 alphabet — `111010111` (weight 7, length 9), `110110101` (weight 6, length 9) and `1011011011` (weight 7, length 10). At query time, seeds are extracted under each pattern, hits into each inverted index are weighted by inverse document frequency, and hit counts are OR-combined across the three patterns. The 500 database chains with the largest accumulated score pass through to exhaustive NW + SW rescoring. In a 200-query evaluation on the 15,391-chain production database, the production prefilter reaches candidate-generation recall@10 = 0.645 and recall@50 = 0.650 at 4.5 ms per query. Alternative designs evaluated and ruled out include RS-20-alphabet variants of the same seed patterns, profile-based and hybrid profile–seed prefilters, and BLOSUM-style neighbourhood seed extension (Supplementary Methods § S3).

### Benchmarks

*The main text reports three complementary benchmarks: hard-negative family classification, family-level retrieval, and geometric fidelity against US-align TM-score*.

#### Hard-negative classification benchmark (Fig. 3a, c)

8,170 same-Rfam-family positive pairs and 3,000 length-matched different-family negative pairs (11,170 total, 102 Rfam families). Negative pairs are matched to the positive length distribution to prevent length-based shortcuts. Performance is measured by ROC AUC on the composite NW+SW score.

#### Retrieval benchmark (Fig. 3b)

156 query chains drawn from 78 Rfam families (2 queries per family) are searched against a 1,041-chain database (≤ 20 chains per family). Recall@*K* is reported for *K* □ {1, 5, 10, 20, 50}.

### Baselines

*We compare RiboSeek with published RNA structural alphabets, a sequence-based BLAST baseline and an RNA-adapted Foldseek control. Where the original code or server is no longer available, legacy methods are reimplemented within our common scoring-and-alignment framework to improve comparability*.

#### iPARTS and iPARTS2

We reimplement iPARTS[6] (η, θ + 23 AP centres) and iPARTS2[7] (η, θ + base type, 92 AP centres) by loading the official cluster centres published in ref.[9] Table 1 and running the same C-accelerated Needleman–Wunsch / Smith–Waterman aligner used throughout this work. Hard-negative benchmark AUCs are reported in the main text (Fig. 3a).

#### R3D-BLAST2

Same 23-letter alphabet as iPARTS. The reference pipeline uses NCBI BLAST on the encoded FASTA; we substitute our C-accelerated NW/SW aligner with the same substitution matrix. Hard-negative AUC is reported in the main text (Fig. 3a).

#### NCBI BLAST bit-score

Sequence-only baseline using `blastn`, word size 7, e-value ≤ 10; bit-score is the discriminant.

#### k6 Jaccard (Supplementary)

Legacy 6-mer Jaccard sequence baseline retained only where indel-sensitivity diagnostics are required (Supplementary Methods S1).

#### Foldseek pseudo-protein control

To assess how well an encoder trained on protein geometry transfers to RNA, we converted each RNA chain to a pseudo-protein PDB by atom mapping (P → N, C4’ → Cα, C1’ → C, base centroid → Cβ) and residue mapping (A → Ala, U → Val, C → Cys, G → Gly; three-letter codes), then ran Foldseek[3] `easy-search` all-versus-all at sensitivity 7.5. A four-way residue-mapping sensitivity analysis (mnemonic, size-matched, polarity-matched, random; Supplementary Fig. S4) confirms that the results are not an artefact of the mapping choice.

### US-align ground truth

We computed US-align[11] TM-score on 600 RNA chain pairs stratified by length (medium 30–100 nt, long 100–500 nt, very long 500+ nt) for Fig. 3c and Fig. 3d, and on 100 dark-family top hits (Fig. 5f) + 27 hub-validation queries (Fig. 5d) for the predicted-library analysis.

### RhoFold+ prediction pipeline

RhoFold+[2] was run on an 8 × A100 80 GB server. For the dark-family search, predictions used single-sequence mode (no MSA input, no Amber relaxation) with six parallel GPU workers, producing approximately 15 predictions per minute. For the MSA-mode hub validation, the Rfam.seed Stockholm alignment for each family was converted to a3m format and passed as the input MSA.

Predicted PDB files were parsed with the same feature-extraction pipeline used for experimental structures and encoded with the production K-means centroids. Chains that failed prediction or yielded fewer than 10 valid RS-20 letters after encoding were discarded; 3,680 of the 3,694 attempted predictions (99.6 %) were retained for downstream analysis.

### Dark-family selection and search

From Rfam.seed (v14.10, 4,227 families), we excluded 157 families represented in our experimental dataset, leaving 4,070 dark families. Of these, 3,694 had ≥ 1 representative sequence of length 40–200 nt containing only AUGC bases; one median-length seed per family was chosen. After RhoFold+ inference and feature filtering, 3,680 predictions were retained and searched against the full 15,391-chain experimental database using 16-worker multi-process parallelized NW+SW scoring. The top 100 dark queries ranked by RS-80 top-1 score were then subjected to US-align verification against their highest-scoring experimental chain.

### Dark-hub validation

For each of the five identified hubs (≥ 3 converging dark families), we sampled additional seeds from Rfam.seed disjoint from the original representative: an initial round of 2–3 seeds per member family (27 seeds) followed by an extension round of up to 6 per family (21 additional seeds), totalling 48 independent validation queries across 19 dark families. Each seed was predicted with RhoFold+, encoded, searched and US-aligned against the expected hub chain. Top-1 hub recall is aggregated weighted-by-family. To test robustness to prediction quality, the initial 27 seeds were also re-predicted in MSA mode using each family’s Rfam.seed Stockholm alignment converted to a3m format as the input MSA — a high-quality hand-curated alignment that avoids the infrastructure cost of an rMSA + RNAcentral pipeline.

### Dark-hub robustness control

To rule out a RhoFold+ fold-attractor bias, we applied an identical pipeline to four control Rfam families with known distinct folds (RF00001 5S rRNA, RF00005 tRNA, RF00162 SAM-I riboswitch, RF00167 purine riboswitch), three independent seeds each (12 total). Aggregate top-1 concentration — the largest fraction of control seeds landing on any single experimental chain — was compared to the dark hubs’ weighted top-1 recall.

### Rfam annotation cross-check

Hub annotation status was cross-validated against the current Rfam 15.x PDB mapping (`pdb_full_region.txt.gz`, April 2026 FTP release) to exclude the possibility that the “unannotated” status of 4 of 5 hubs reflects stale mapping data.

### Statistical analysis

Bootstrap confidence intervals on AUC used 1,000 resamples with replacement from the full pair list; reported CIs are the empirical 2.5th–97.5th percentile.

### Learned prefilter (exploratory)

Exploratory learned-prefilter experiments are described in Supplementary Methods § S8 but are not part of the production method evaluated in the main text.

## Data and code availability

Code for RiboSeek, including the C-accelerated NW/SW aligner, trained RS-20 and RS-80 centroids, all substitution matrices, the spaced-seed prefilter, feature-extraction scripts, and baseline reproduction code, is available at https://github.com/void-echo/RiboSeek. The repository will be made fully public upon journal publication; in the interim, access is available from the corresponding authors upon reasonable request. Processed files can be reproduced from publicly available RCSB mmCIF files (rcsb.org) and Rfam annotations (rfam.org) using the included scripts. US-align was obtained from the Zhang lab (zhanggroup.org).

## Author contributions

D.W. conceived the study, developed the method, performed the analyses and drafted the manuscript. J.J. contributed to benchmark construction and baseline re-implementation. J.Q. contributed to data curation and manuscript revision. L.W. advised on methodological framing and benchmark scope. S.W. and Q.L. supervised the study, secured resources and revised the manuscript. All authors approved the final version.

## Acknowledgements

The authors thank colleagues for helpful discussions.

## References

1. Abramson, J. et al. Accurate structure prediction of biomolecular interactions with AlphaFold 3. Nature 630, 493–500 (2024).

2. Shen, T. et al. Accurate RNA 3D structure prediction using a language model-based deep learning approach. Nat. Methods 21, 2287–2298 (2024).

3. van Kempen, M. et al. Fast and accurate protein structure search with Foldseek. Nat. Biotechnol. 42, 243–246 (2024).

4. Barrio-Hernandez, I. et al. Clustering-predicted structures at the scale of the known protein universe. Nature 622, 637–645 (2023).

5. Chang, Y.-F., Huang, Y.-L. & Lu, C. L. SARSA: a web tool for structural alignment of RNA using a structural alphabet. Nucleic Acids Res. 36, W19–W24 (2008).

6. Wang, C.-W., Chen, K.-T. & Lu, C. L. iPARTS: an improved tool of pairwise alignment of RNA tertiary structures. Nucleic Acids Res. 38, W340–W347 (2010).

7. Yang, C.-H. et al. iPARTS2: an improved tool for pairwise alignment of RNA tertiary structures, version 2. Nucleic Acids Res. 44, W328–W332 (2016).

8. Liu, Y.-C. et al. R3D-BLAST: a search tool for similar RNA 3D substructures. Nucleic Acids Res. 39, W45–W49 (2011).

9. Yen, C.-Y. et al. R3D-BLAST2: an improved BLAST-based tool for searching similar RNA 3D substructures. Nucleic Acids Res. 45, W129–W134 (2017).

10. Ontiveros-Palacios, N. et al. Rfam 15: RNA families database in 2025. Nucleic Acids Res. 53, D258–D267 (2025).

11. Zhang, C., Shine, M., Pyle, A. M. & Zhang, Y. US-align: universal structure alignments of proteins, nucleic acids, and macromolecular complexes. Nat. Methods 19, 1109–1115 (2022).

12. Altschul, S. F., Gish, W., Miller, W., Myers, E. W. & Lipman, D. J. Basic local alignment search tool. J. Mol. Biol. 215, 403–410 (1990).

13. Baek, M. et al. Accurate prediction of protein-nucleic acid complexes using RoseTTAFold2NA. Nat. Methods 21, 117–121 (2024).

14. Needleman, S. B. & Wunsch, C. D. A general method applicable to the search for similarities in the amino acid sequence of two proteins. J. Mol. Biol. 48, 443–453 (1970).

15. Smith, T. F. & Waterman, M. S. Identification of common molecular subsequences. J. Mol. Biol. 147, 195–197 (1981).

16. Nawrocki, E. P. & Eddy, S. R. Infernal 1.1: 100-fold faster RNA homology searches. Bioinformatics 29, 2933–2935 (2013).

17. Leontis, N. B., Lescoute, A. & Westhof, E. The building blocks and motifs of RNA architecture. Curr. Opin. Struct. Biol. 16, 279–287 (2006).

18. Čech, P., Svozil, D. & Schneider, B. DNATCO: assignment of DNA conformers at dnatco.org. Nucleic Acids Res. 41, W284–W287 (2013).

19. Wadley, L. M., Keating, K. S., Duarte, C. M. & Pyle, A. M. Evaluating and learning from RNA pseudotorsional space: quantitative validation of a reduced representation for RNA structure. J. Mol. Biol. 372, 942–957 (2007).

20. Duarte, C. M. & Pyle, A. M. Stepping through an RNA structure: a novel approach to conformational analysis. J. Mol. Biol. 284, 1465–1478 (1998).

21. Jumper, J. et al. Highly accurate protein structure prediction with AlphaFold. Nature 596, 583–589 (2021).

22. Steinegger, M. & Söding, J. MMseqs2 enables sensitive protein sequence searching for the analysis of massive data sets. Nat. Biotechnol. 35, 1026–1028 (2017).

23. Griffiths-Jones, S. miRBase: the microRNA sequence database. Methods Mol. Biol. 342, 129–138 (2006).

24. Nawrocki, E. P. Annotating functional RNAs in genomes using Infernal. Methods Mol. Biol. 1097, 163–197 (2014).

25. Parisien, M. & Major, F. The MC-Fold and MC-Sym pipeline infers RNA structure from sequence data. Nature 452, 51–55 (2008).

26. Sarver, M., Zirbel, C. L., Stombaugh, J., Mokdad, A. & Leontis, N. B. FR3D: finding local and composite recurrent structural motifs in RNA 3D structures. J. Math. Biol. 56, 215–252 (2008).

27. Petrov, A. I., Zirbel, C. L. & Leontis, N. B. Automated classification of RNA 3D motifs and the RNA 3D Motif Atlas. RNA 19, 1327–1340 (2013).

28. Schrodinger, LLC. The PyMOL Molecular Graphics System, Version 2.5. (2021). [http://www.pymol.org]

29. Burley, S. K. et al. RCSB Protein Data Bank: celebrating 50 years of the PDB with new tools for understanding and visualizing biological macromolecules. Protein Sci. 31, 187–208 (2022).

30. Waldispühl, J. & Reinharz, V. RNA structure motifs: a neural network perspective. Curr. Opin. Struct. Biol. 76, 102369 (2022).

31. Su, J. et al. SaProt: protein language modeling with structure-aware vocabulary. International Conference on Learning Representations (ICLR) (2024).

32. Cormack, G. V., Clarke, C. L. A. & Büttcher, S. Reciprocal rank fusion outperforms condorcet and individual rank learning methods. In Proceedings of the 32nd International ACM SIGIR Conference on Research and Development in Information Retrieval, 758–759 (2009).

33. Chen, K. et al. OmniGenome: aligning RNA sequences with secondary structures in foundation models. Preprint at arXiv 2407.11242 (2024).

34. Penić, R. J. et al. RiNALMo: general-purpose RNA language models can generalize well on structure prediction tasks. Preprint at arXiv 2403.00043 (2024).

35. Chen, J. et al. Interpretable RNA foundation model from unannotated data for highly accurate RNA structure and function predictions. arXiv 2204.00300 (2022). [RNA-FM]

36. Yin, W. et al. ERNIE-RNA: an RNA language model with structure-enhanced representations. Preprint at bioRxiv 2024.03.17.585376 (2024).

