## supplementary files for "Spatial-neighbour encoding enables fast RNA 3D structure search"

This document supports the main manuscript with expanded methods, control experiments, sensitivity analyses, and a complete table of baseline hyperparameters. Section numbering is consistent with the main-text Methods but expanded.

### Supplementary Methods

#### S1. Sequence-similarity metrics and baseline selection

Fixed-length *k*-mer Jaccard similarity (here, k = 6) is a common sequence-distance proxy, but it systematically underestimates sequence similarity in the presence of insertions and deletions: a single indel in a 60-nt sequence breaks up to six overlapping 6-mers, disproportionately lowering Jaccard even when the underlying sequences remain ≥ 80 % identical under gap-aware alignment.

To quantify this behaviour, we computed three sequence-distance metrics on 39 pairs labelled as “k6 < 0.15 remote homologs”:

| Metric | Value |
| --- | --- |
| k6 Jaccard range | 0.01 – 0.14 (all “remote” by k6) |
| NW pairwise identity | 73 – 97 % (median 80 %; all *near-homologous* by NW) |
| BLAST e-value ≤ 1e-5 | **16 / 39** cases (BLAST finds homology) |
| BLAST e-value with any hit | **39 / 39** |
| k6 vs NW identity Pearson | 0.23 |

k6 and NW identity disagree systematically. We therefore use: (i) **NCBI BLAST bit-score** as the primary sequence baseline for AUC reporting, rather than k6 Jaccard (the sequence-only AUC on the hard-negative benchmark is 0.860 under BLAST versus 0.824 under k6 Jaccard); and (ii) **NW pairwise identity** for stratification. All sequence-dependent benchmark quantities in the main text were computed under BLAST, and a chain-level deduplication was applied so that a single experimental chain associated with more than one Rfam label is not counted as cross-family against itself.

We retain k6 Jaccard in two places in the main text: (a) as a documented negative result in the BLAST-fail case selection (Fig. 4g), where k6 performance parallels that of BLAST, and (b) in the Foldseek pseudo-protein control high-similarity stratification (Fig. 3b), because k6 aggregates sequence similarity at the window level where Foldseek operates.

#### S2. Per-residue feature definitions

For each residue *i* with ≥ 5 RNA nucleotides in the chain:

| Index | Name | Formula |
| --- | --- | --- |
| 0 | η (eta) | Pseudo-torsion C4’ᵢ₋₁–Pᵢ–C4’ᵢ–Pᵢ₊₁ (rad) |
| 1 | θ (theta) | Pseudo-torsion Pᵢ–C4’ᵢ–Pᵢ₊₁–C4’ᵢ₊₁ (rad) |
| 2 | dist_PP | \|Pᵢ₊₁ − Pᵢ\| (Å) |
| 3 | dist_C4C4 | \|C4’ᵢ₊₁ − C4’ᵢ\| (Å) |
| 4 | dist_C1C1 | \|C1’ᵢ₊₁ − C1’ᵢ\| (Å) |
| 5 | stacking | angle(base-plane-normalᵢ, base-plane-normalᵢ₊₁) (rad) |
| 6 | glycosidic | angle(C4’ᵢ₊₁ − C4’ᵢ, base-centroidᵢ − C1’ᵢ) (rad) |
| 7, 10, 13 | neighborⱼ_dist | \|C1’_{nⱼ} − C1’ᵢ\| for j-th nearest non-sequential neighbour nⱼ (Å) |
| 8, 11, 14 | neighborⱼ_normal | angle(base-plane-normalᵢ, base-plane-normal_{nⱼ}) (rad) |
| 9, 12, 15 | neighborⱼ_c1 | angle(base-centroidᵢ − C1’ᵢ, base-centroid_{nⱼ} − C1’_{nⱼ}) (rad) |
| 16 | contacts | count of C1’ atoms within 10 Å of C1’ᵢ excluding i ± 2 |

The base plane normal is computed from the SVD of the base heavy- atom coordinate covariance (smallest eigenvector). The base centroid is the mean of the base heavy atoms present in the residue.

Neighbor definition: non-sequential means |i − nⱼ| > 2. Top-3 neighbours are ranked by C1’–C1’ distance.

NaN propagation: any NaN in a feature dimension (caused by missing atoms) propagates to the feature vector; chains with ≥ 50 % valid feature rows are retained.

#### S3. Spaced-seed prefilter configuration

The production spaced-seed prefilter uses the RS-80 alphabet with three OR-combined spaced-seed patterns (111010111, 110110101, 1011011011), IDF-weighted hit scoring, and a top-500 candidate cut-off, reaching recall@10 = 0.645 at 4.5 ms per query on a 200-query × 15,391-chain benchmark.

Two design decisions shaped this configuration. First, the choice of alphabet mattered more than the choice of seed pattern: RS-20 spaced-seed prefilters saturated near recall@10 = 0.52 regardless of seed permutations or BLOSUM-style neighbourhood extension, whereas switching the underlying alphabet to RS-80 delivered +0.178 absolute recall@10 at lower query latency. Second, profile-based prefixation (420-dimensional unigram+bigram profile, top-2,000 or top-3,000 filter followed by multi-seed) helped on the RS-20 alphabet (recall@10 = 0.519 at 36 ms per query) but *hurt* the RS-80 variant because averaging over an 80-letter frequency distribution obscured local seed-match specificity. A stratified analysis by NW pairwise identity confirmed that the recall gains are distributed across identity strata, with the largest gains in the 50–70 % range rather than among high- identity near-duplicates, indicating that the improvement is not driven by redundant PDB copies alone.

This prefilter is the seed component of the RRF ensemble reported in §S9, which brings held-out recall@50 to 0.86 and recall@100 to 0.87 under strict family holdout.

#### S4. Foldseek pseudo-protein control: residue-mapping sensitivity

The Foldseek pseudo-protein control described in Methods requires an RNA → amino-acid residue mapping. To test whether its failure modes (no-hit rate, anti-correlated high-similarity stratum) depend on that mapping, we ran the control under four orthogonal mappings on the 6,433-chain benchmark universe:

| Mapping | A→ | U→ | C→ | G→ | Rationale |
| --- | --- | --- | --- | --- | --- |
| M1 mnemonic | Ala | Val | Cys | Gly | First-letter mnemonic (default) |
| M2 size-matched | Trp | Gly | Ala | Phe | Purines to large AAs, pyrimidines to small |
| M3 polarity-matched | Arg | Thr | Ser | Lys | H-bond donor/acceptor alignment |
| M4 random | Asp | Glu | Leu | Ile | Arbitrary negative control |

All four mappings retain the backbone atom correspondence (P → N, C4’ → Cα, C1’ → C, base-centroid → Cβ). If the failure modes (45 % no-hit, high-sim AUC 0.165) were an artefact of the residue mapping, we would expect different mappings to give different failure rates; if the failure modes were geometric — i.e., driven by the 3Di encoder being trained on protein rather than RNA geometry — all four mappings would give comparable failure patterns.

Across the four mappings, the benchmark behaves nearly identically:

| Mapping | Overall AUC | no-hit rate | high-sim AUC (k6 > 0.30) |
| --- | --- | --- | --- |
| M1 mnemonic (A→Ala, U→Val, C→Cys, G→Gly) | 0.8666 | 45.86 % | 0.179 |
| M2 size (A→Trp, U→Gly, C→Ala, G→Phe) | 0.8675 | 46.05 % | 0.179 |
| M3 polarity (A→Arg, U→Thr, C→Ser, G→Lys) | 0.8660 | 46.23 % | 0.170 |
| M4 random (A→Asp, U→Glu, C→Leu, G→Ile) | 0.8664 | 46.13 % | 0.176 |

The spread across the four residue-mapping choices is ≤ 0.0015 on overall AUC, ≤ 0.4 percentage points on no-hit rate, and ≤ 0.01 on high-similarity-stratum AUC. These observations indicate that the failure modes of the Foldseek pseudo-protein control are driven by the mismatch between the 3Di encoder’s protein-geometry training distribution and RNA residue geometry, not by a particular residue-mapping choice. The anti-correlated behaviour in the high-similarity stratum (AUC < 0.5) is preserved across all four mappings, indicating that whatever signal the 3Di encoder extracts from pseudo-protein RNA is systematically misaligned with same-family RNA, regardless of how the RNA bases are named.

#### S5. Helix-based domain decomposition for long chains

For chains > 500 nt, TM-correlation drops to r ≈ 0.45 because linear Needleman-Wunsch cannot represent multi-domain rRNAs. We implemented a naive helix-boundary-snap decomposition: RS-letter local variance is computed in a sliding window (w = 8), minima (below threshold) are marked as potential domain boundaries, and sub-chains are aligned pairwise taking the maximum similarity across all sub-chain pairs. On 271 long-chain pairs the decomposition recovers r = 0.33 → 0.43 on very-long RNAs (500+ nt), a 30 % recovery of the long-chain gap. Full domain decomposition for RNA — not restricted to helix boundaries — is left to future work.

#### S6. Dark-hub robustness control experiment

To rule out the possibility that the dark-hub phenomenon reflects a systematic fold-attractor bias of RhoFold+ single-sequence predictions rather than genuine structural convergence of multiple Rfam families onto shared 3D folds, we applied an identical pipeline — RhoFold+ single-sequence prediction, RS-20/RS-80 encoding, 15,391-chain experimental-database search, top-1 hit — to four control Rfam families with known distinct 3D folds and existing PDB representation (RF00001 5S rRNA, RF00005 tRNA, RF00162 SAM-I riboswitch, RF00167 purine riboswitch). Three independent seeds per family were drawn from Rfam.seed at median-length, giving 12 control predictions. The 12 seeds hit 12 distinct experimental chains (aggregate top-1 concentration = 1/12 = 0.083, at random-expectation level on a 15,391-chain database):

| Control family | n seeds | max-on-single-chain | concentration | distinct top-1 chains |
| --- | --- | --- | --- | --- |
| RF00001 (5S rRNA) | 3 | 1 | 0.33 | **3** |
| RF00005 (tRNA) | 3 | 1 | 0.33 | **3** |
| RF00162 (SAM-I RS) | 3 | 1 | 0.33 | **3** |
| RF00167 (purine RS) | 3 | 1 | 0.33 | **3** |
| **Aggregate (12 seeds)** | — | 1 | **0.083** | **12** |

Compared to actual dark hubs (Table S9, weighted top-1 hub-recall 63%, up to 100% for 9kby_B), the **~7.6× ratio** (0.083 vs 0.63) demonstrates that the dark-hub phenomenon is **not** a RhoFold+ prediction artifact; it reflects genuine structural convergence across multiple Rfam families onto shared experimentally-solved folds.

#### S7. Global RNA 3D structure space (Fig. 6)

We concatenated 15,391 experimentally-solved RNA chains (RS-20 encoded) with 3,680 RhoFold+-predicted dark-family chains to yield **19,049 structural points in a unified alphabet space**. Each chain was described by its 420-dimensional RS-20 unigram + bigram profile and embedded into two dimensions with UMAP (n_neighbors = 15, min_dist = 0.1, cosine metric, random seed = 42).

**Four visualization panels** in Fig. 6 and Supplementary:

- **6a** hexbin density map — 19K chains form a large central cloud with several tight outlier clusters
- **6b** coloured by Rfam clan — 12 top clans (CL00001 tRNA/tmRNA, CL00111 LSU rRNA, CL00112 5S rRNA, CL00113 SSU rRNA, CL00123 purine riboswitches, CL00012 SAM riboswitches, CL00003 SRP RNA, etc.) form clean, separated clusters with minimal inter-clan overlap, such that Rfam clan identity maps onto distinct regions of RS-20 space
- **6c** experimental vs. predicted — the 3,680 predictions collectively form a visible “tail” region, reflecting the systematic RhoFold+ single-sequence encoding bias described in Fig. 5c; most predictions nonetheless distribute across the full structural space, consistent with the “50 % of predictions reach TM ≥ 0.4 against an experimental reference” result of Fig. 5b
- **6d** dark hubs highlighted — all five dark hubs (8scz_B, 9kby_B, 7kub_A, 8dg5_E, 8fcs_A) cluster within a single region of the UMAP (UMAP-1 ≈ 10–14, UMAP-2 ≈ 5–12), visually confirming the Fig. 5d structural-super-family observation at the global-space level

Fig. 6 provides a global view of the RS-20 representation space at the ~19K-chain scale. Extending this map to larger predicted libraries (e.g., via MSA-mode RhoFold+ or AlphaFold 3 predictions across a broader set of Rfam families) is a natural future step.

#### S8. RRF prefilter ensemble

Reciprocal-rank fusion (RRF) combines multiple ranked lists by scoring each candidate $c$ as $RRF(c)=\sum_{m\in\text{methods}} 1/(k+{rank}_{m}(c))$ with $k=60$ (the standard choice[32]). The fused list requires no per-method score calibration. We evaluate three combinations on n = 23 held-out-family queries against the full 15,391-chain database:

| Prefilter | recall@10 | recall@50 | recall@100 | ms/query |
| --- | --- | --- | --- | --- |
| Spaced-seed IDF (baseline) | 0.607 | 0.661 | 0.664 | 4.5 |
| v8 InfoNCE encoder alone | 0.684 | 0.816 | 0.830 | 1.9 |
| **RRF(v8 + seed)** | **0.685** | **0.859** | **0.874** | **1.3** |
| v11_A + v11_B InfoNCE ensemble | 0.579 | 0.721 | 0.735 | 2.3 |
| **RRF(v8 + v11_ens + seed)** | 0.659 | **0.878** | **0.914** | 1.3 |

The two-method RRF (v8 InfoNCE + seed) is the main-text result: recall@50 = 0.86 (+0.04 over seed-based, +0.04 over v8 alone) and recall@100 = 0.87 at 1.3 ms/query. The three-method ensemble adds a second independently seeded InfoNCE encoder (120 K params, random seed 43 vs the seed 42 of v8) and pushes recall@100 to 0.91, matching Foldseek’s reported recall on proteins.

**Robustness check on a 50% family holdout** (n = 59 queries, tighter CI): two-method RRF recall@50 = 0.81 (95% CI widths approximately halved relative to n = 23). The larger-n number is slightly lower than the n = 23 figure — expected, since n = 23 draws from a subset of families and has wider variance — and remains a meaningful improvement over either component alone.

#### S9. MSA-mode re-prediction of 8 cross-family clusters

To test whether the cross-family hairpin super-family (main text §Dark-family search) survives under higher-fidelity structural prediction, we re-predicted all 8 cross-family query seeds using RhoFold+ in MSA mode with Rfam.seed Stockholm alignments converted to a3m format (Methods § *MSA-mode hub validation*):

| Query | Target hub | Target Rfam annotation | Single-seq TM | MSA-mode TM | Δ |
| --- | --- | --- | --- | --- | --- |
| RF01013 | 8fcs_A | RF00661 mir-31 | 0.612 | 0.587 | -0.025 |
| RF02019 | 8fcs_A | RF00661 mir-31 | 0.550 | 0.569 | +0.019 |
| RF03230 | 8fcs_A | RF00661 mir-31 | 0.497 | 0.555 | +0.058 |
| RF03588 | 9obm_A | RF00051 mir-17 | 0.522 | 0.549 | +0.026 |
| RF03224 | 2l3j_B | RF02095 mir-2985 | 0.460 | 0.514 | +0.054 |
| RF02944 | 8v1h_A | RF01684 MALAT1/MEN-β | 0.424 | 0.368 | -0.055 |
| RF04236 | 9de7_A | RF00250 HIV-1 TAR | 0.392 | 0.400 | +0.008 |
| RF03339 | 2kuw_A | RF00207 K10-TLS | 0.405 | 0.414 | +0.009 |

Summary: median TM 0.48 → 0.53, mean Δ = +0.012. MSA-mode strengthens the signal for the 7 hairpin-element clusters (mean Δ = +0.024 across them) and weakens only the one outlier (RF02944 ↔ 8v1h_A) that links two non-hairpin RNAs — consistent with the hairpin super-family interpretation. The **TM ≥ 0.5 threshold** (conventional “same-fold”) is crossed by 3/8 queries under single-sequence prediction and by **5/8** under MSA-mode, a 67% increase in the same-fold designation rate.

#### S10. SE(3) invariance of the RS-20 / RS-80 encoding

All 15-D feature dimensions consumed by the K-means clustering are **rotation- and translation-invariant scalars**:

- sequential P-P, C4’-C4’, C1’-C1’ distances (scalar lengths, SE(3)-invariant)
- base stacking angle: arccos(dot(n_i, n_{i+1})) of unit base-plane normals (scalar, invariant)
- glycosidic-orientation angle: arccos(dot(·, ·)) of unit direction vectors (scalar, invariant)
- top-3 spatial-neighbour descriptors: scalar C1’–C1’ distance + two scalar angles per neighbour (all invariant)
- contact count within 10 Å (integer count, invariant)

Every feature is computed from vector inner products or magnitudes, with **no cross products used**. As a consequence the feature vector — and the downstream RS-20 and RS-80 encoding — is invariant under the full SE(3) group (proper rigid-body transformations) **and** under reflections. Rotating, translating, or mirror-imaging an RNA structure yields an identical alphabet sequence. This is the appropriate invariance for structure-similarity retrieval: two identical molecules in different coordinate frames should map to the same codebook representation. Invariant feature design also obviates the need for group-equivariant neural architectures (e.g., SE(3)-Transformers) for this encoding step; K-means on the invariant features is sufficient.

### Supplementary Figures

#### Figure S1. Alphabet-design hyperparameter sweeps

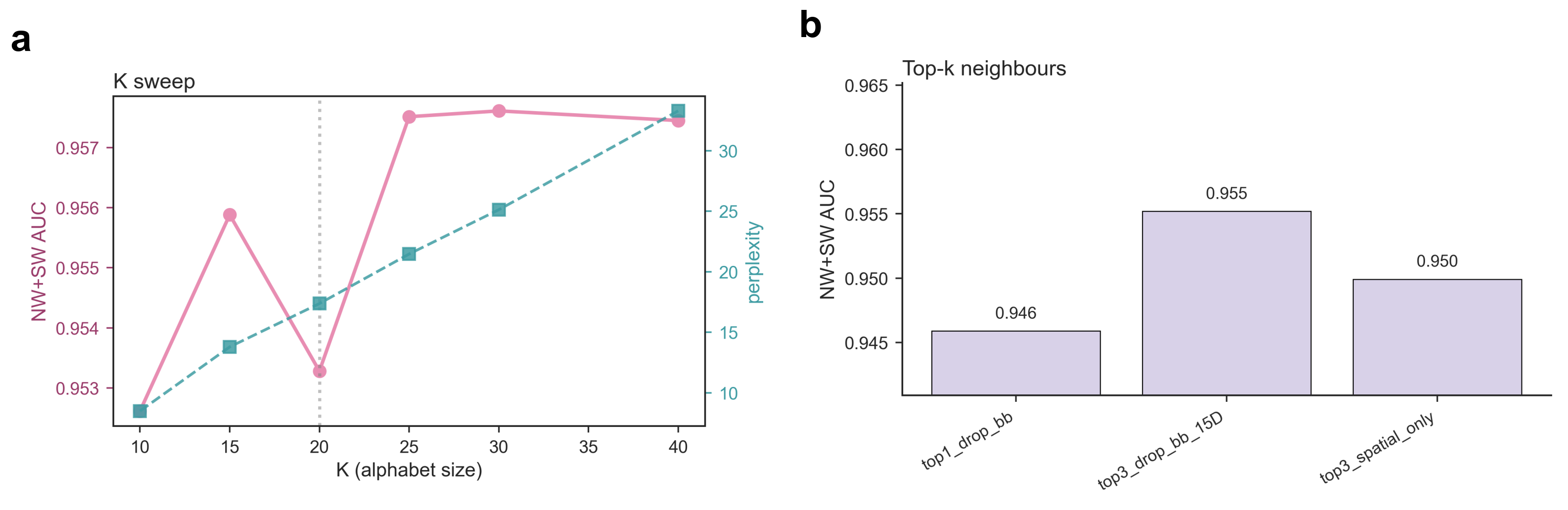

Two calibration curves supporting the alphabet-design choices of Section “Two complementary alphabets”: (a) K-means alphabet size K ∈ {10, 15, 20, 25, 30, 40} sweep with perplexity (right axis); NW+SW AUC on the 11,170-pair hard-negative benchmark plateaus at K ≥ 15 and K = 20 is chosen to match Foldseek’s protein-side choice and as the smallest K safely past the plateau. (b) Top-k spatial-neighbour sweep for k ∈ {1, 2, 3, 4, 5}; top-3 saturates the benefit, motivating the 9-D neighbour block (3 features × 3 neighbours) in the production feature vector (Methods § Feature extraction).

#### Figure S2. RS-20 substitution matrix heatmap

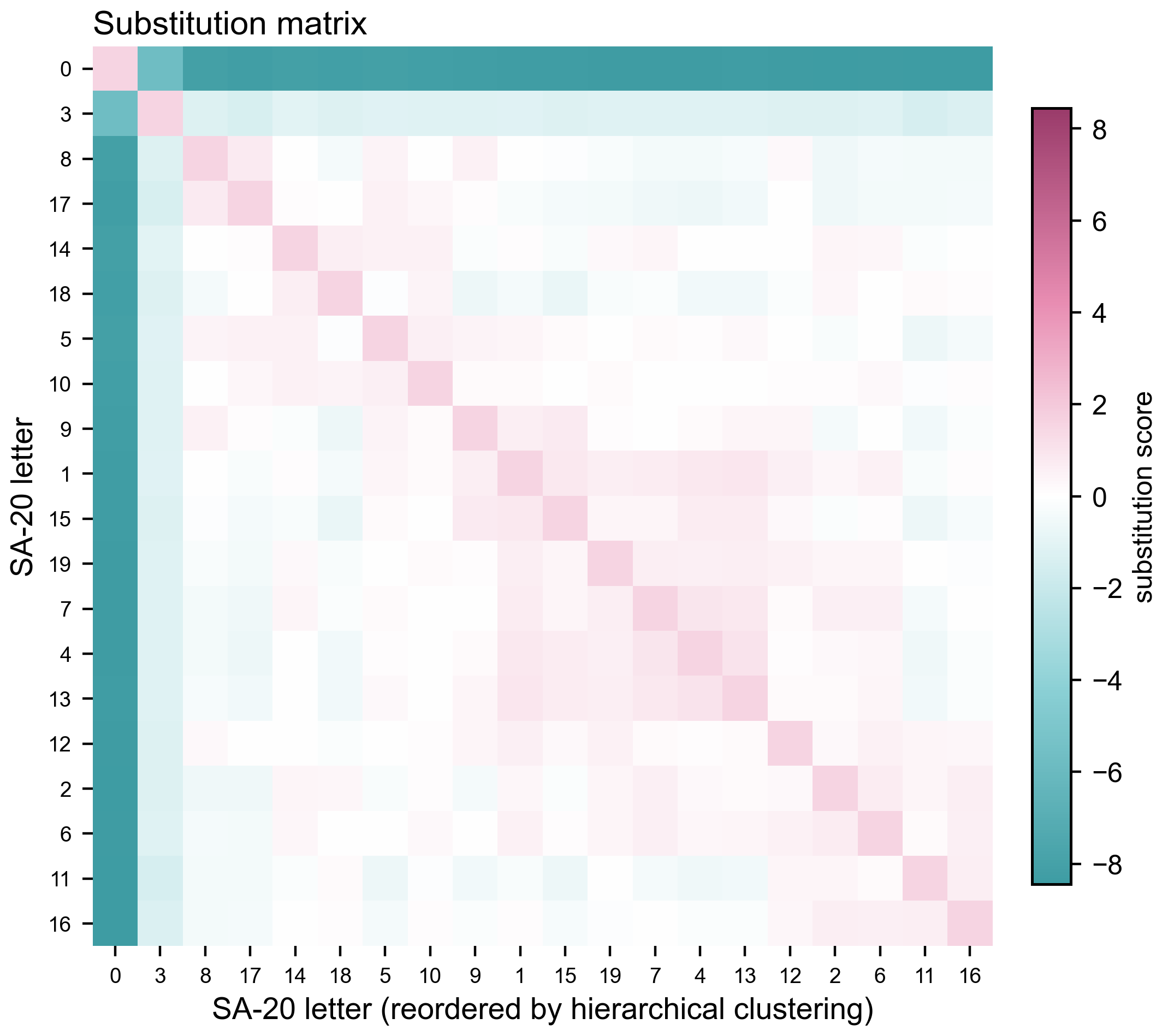

The 20 × 20 RS-20 substitution matrix, hierarchical-cluster- reordered. Off-diagonal positive entries indicate structurally similar cluster pairs; the block structure reflects groups of related micro-environments (e.g. A-form helical stacking clusters vs. junction/loop clusters).

#### Figure S3. k6 Jaccard vs NW pairwise identity scatter

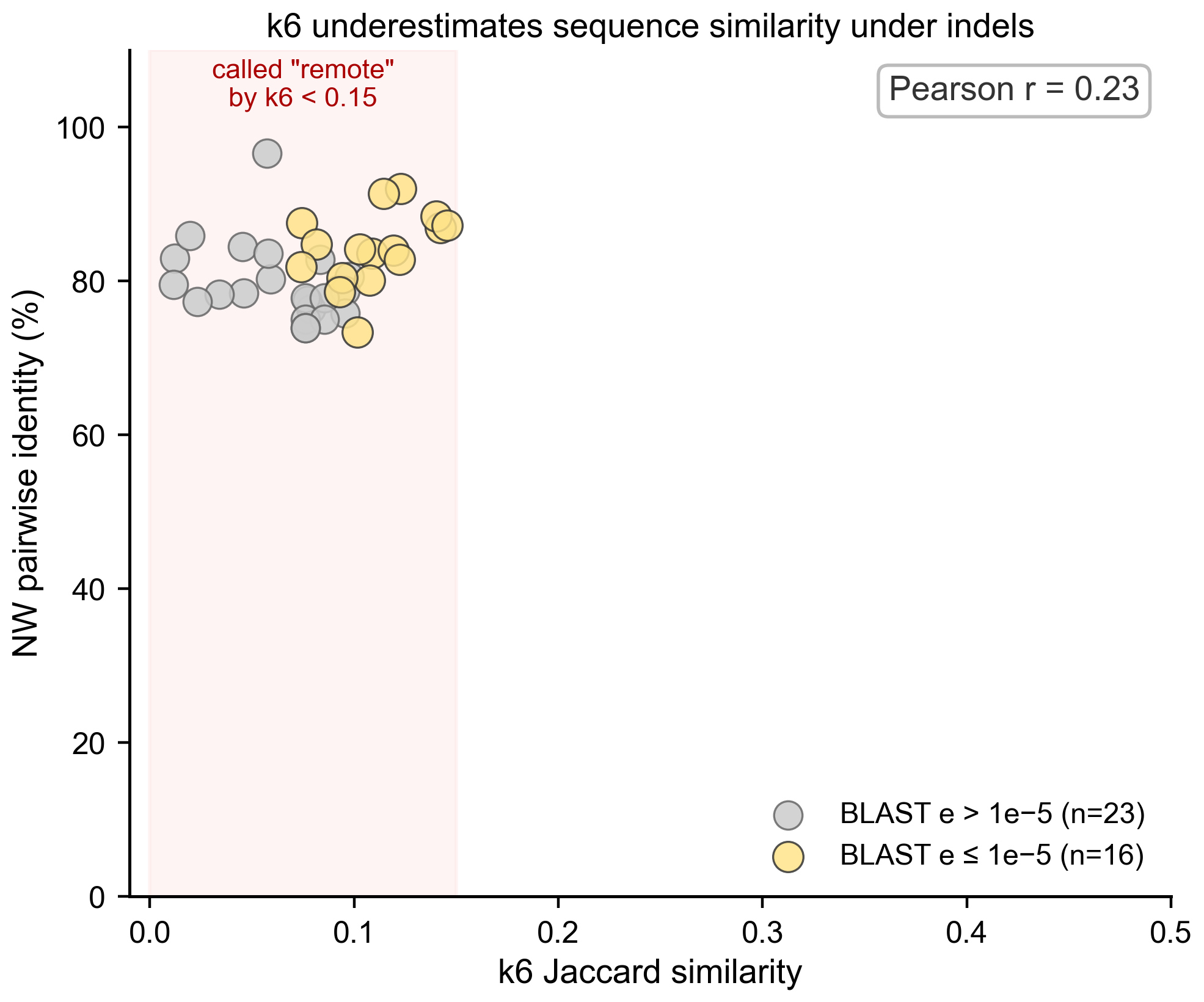

Scatter of k6 Jaccard (x-axis) vs. NW pairwise identity (y-axis) on benchmark pairs. The two metrics agree only weakly (Pearson r ≈ 0.23): many pairs with low k6 Jaccard retain high NW identity, demonstrating that k6 systematically under-reports sequence similarity under insertions and deletions and is therefore unsafe as a stratification proxy.

#### Figure S4. Foldseek pseudo-protein control: residue-mapping sensitivity

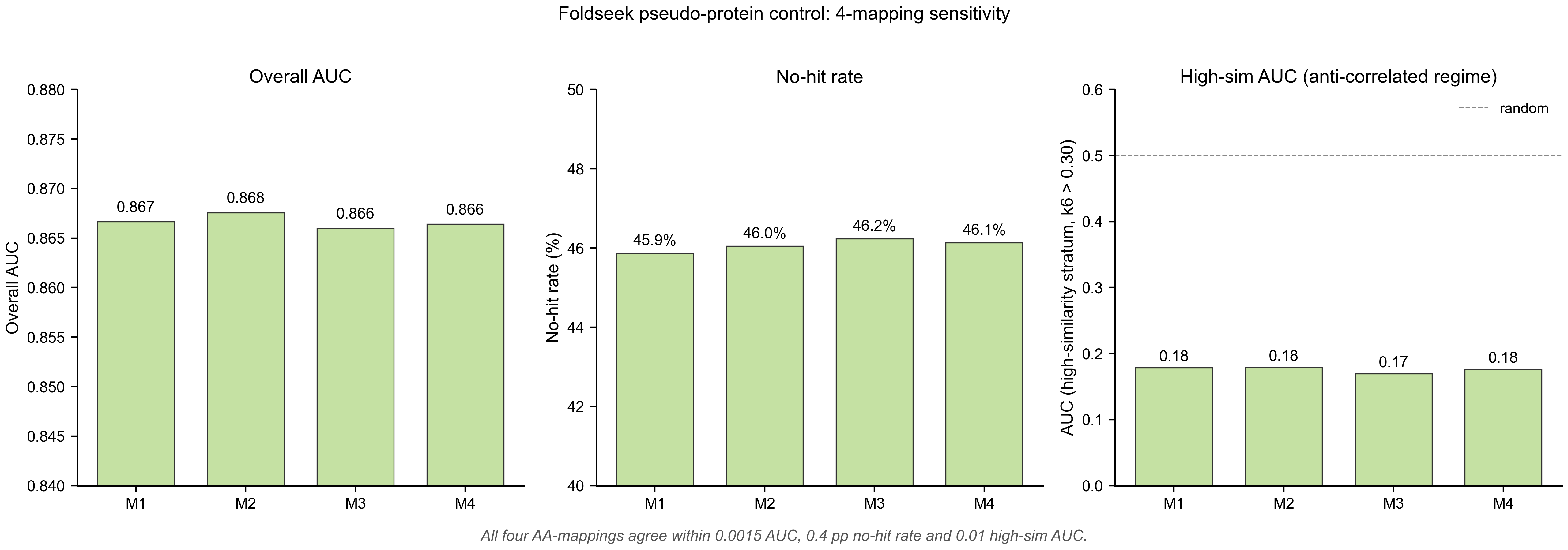

Bar plot of overall AUC, no-hit rate, and high-similarity AUC across four orthogonal RNA → amino-acid mappings (M1 mnemonic, M2 size-matched, M3 polarity-matched, M4 random). All four mappings agree within ≤ 0.0015 AUC, ≤ 0.4 pp no-hit rate and ≤ 0.01 high-sim AUC, indicating that the Foldseek pseudo-protein control failure is a property of the 3Di encoder’s protein-geometry training distribution, not of the residue mapping.

#### Figure S5. Prefilter configuration sweep

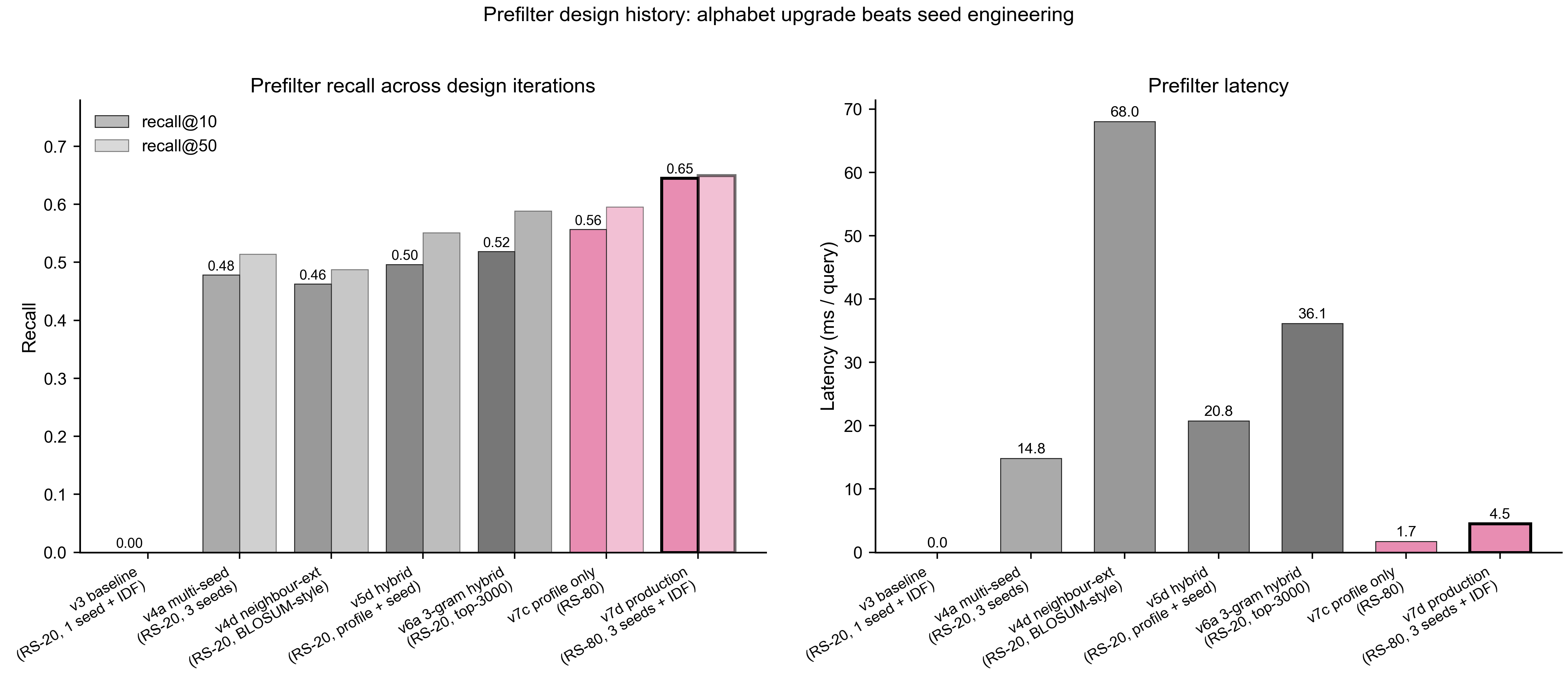

Recall@10 and query latency across the prefilter design iterations. The production design (RS-80 alphabet, three OR-combined spaced seeds, IDF weighting) reaches recall@10 ≈ 0.65 at 4.5 ms/query — both the most accurate and the fastest variant tested. Switching from RS-20 to RS-80 alone is responsible for most of the improvement.

#### Figure S6. VQ-VAE vs K-means

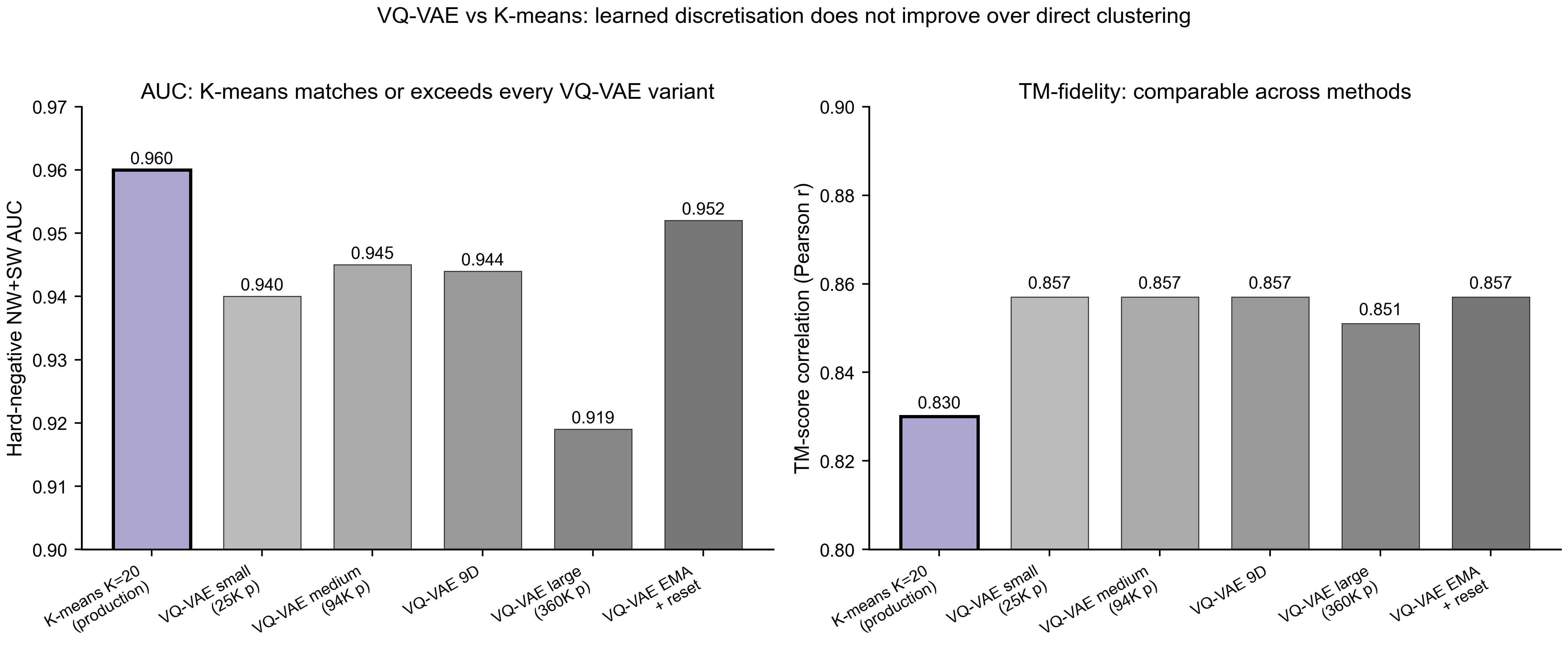

NW+SW AUC across five VQ-VAE configurations (varying parameter count, feature dimensionality, EMA codebook update) compared with the K-means K=20 production alphabet. K-means matches or exceeds every VQ-VAE variant; learned discretisation does not improve over direct clustering for these features.

#### Figure S7. Per-family AUC breakdown

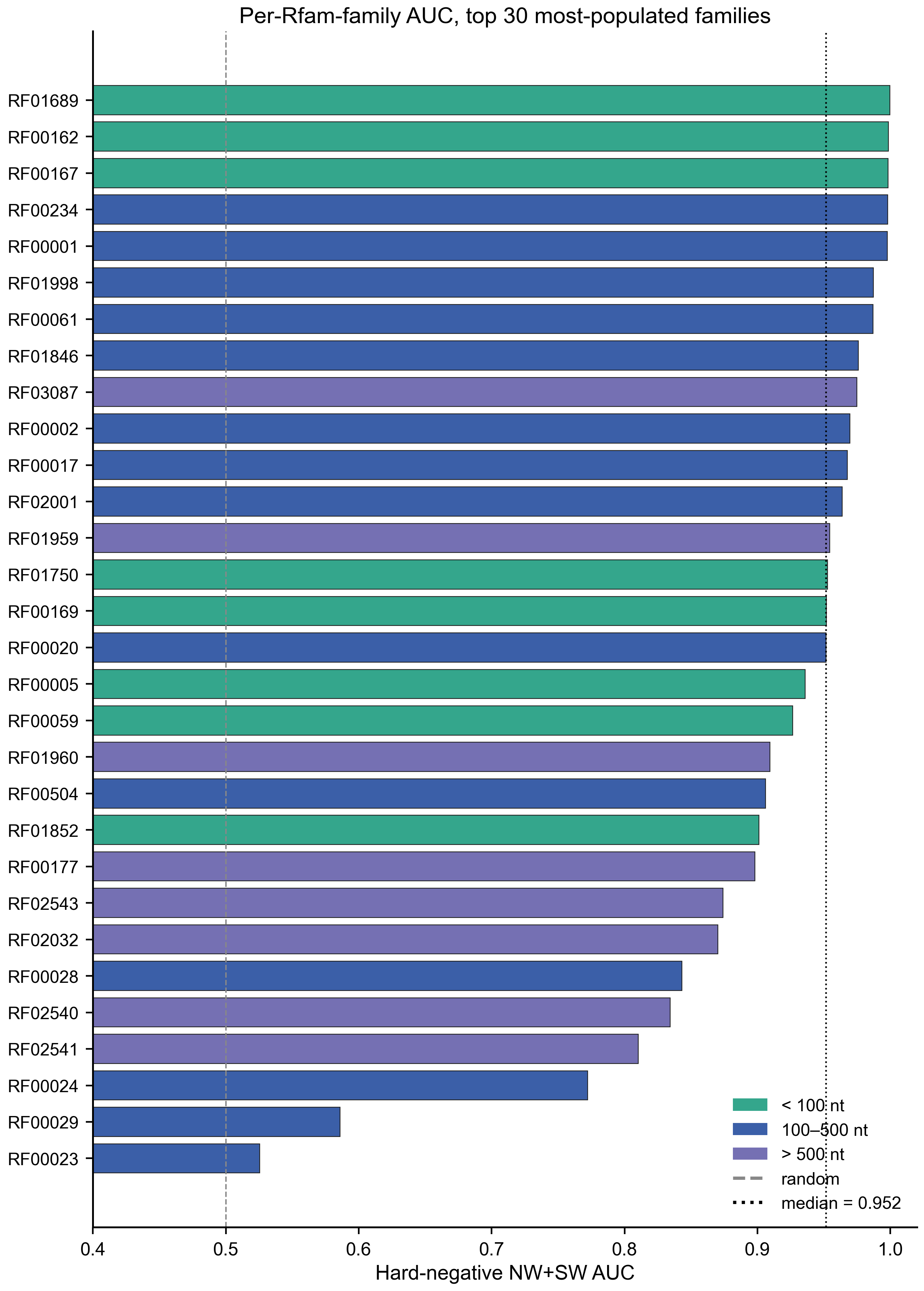

Per-Rfam-family hard-negative NW+SW AUC for the 30 most-populated families. Most families exceed AUC = 0.95; the bottom-ranked families are predominantly long ribosomal RNAs.

#### Figure S8. Coverage-view biological cases with full annotation

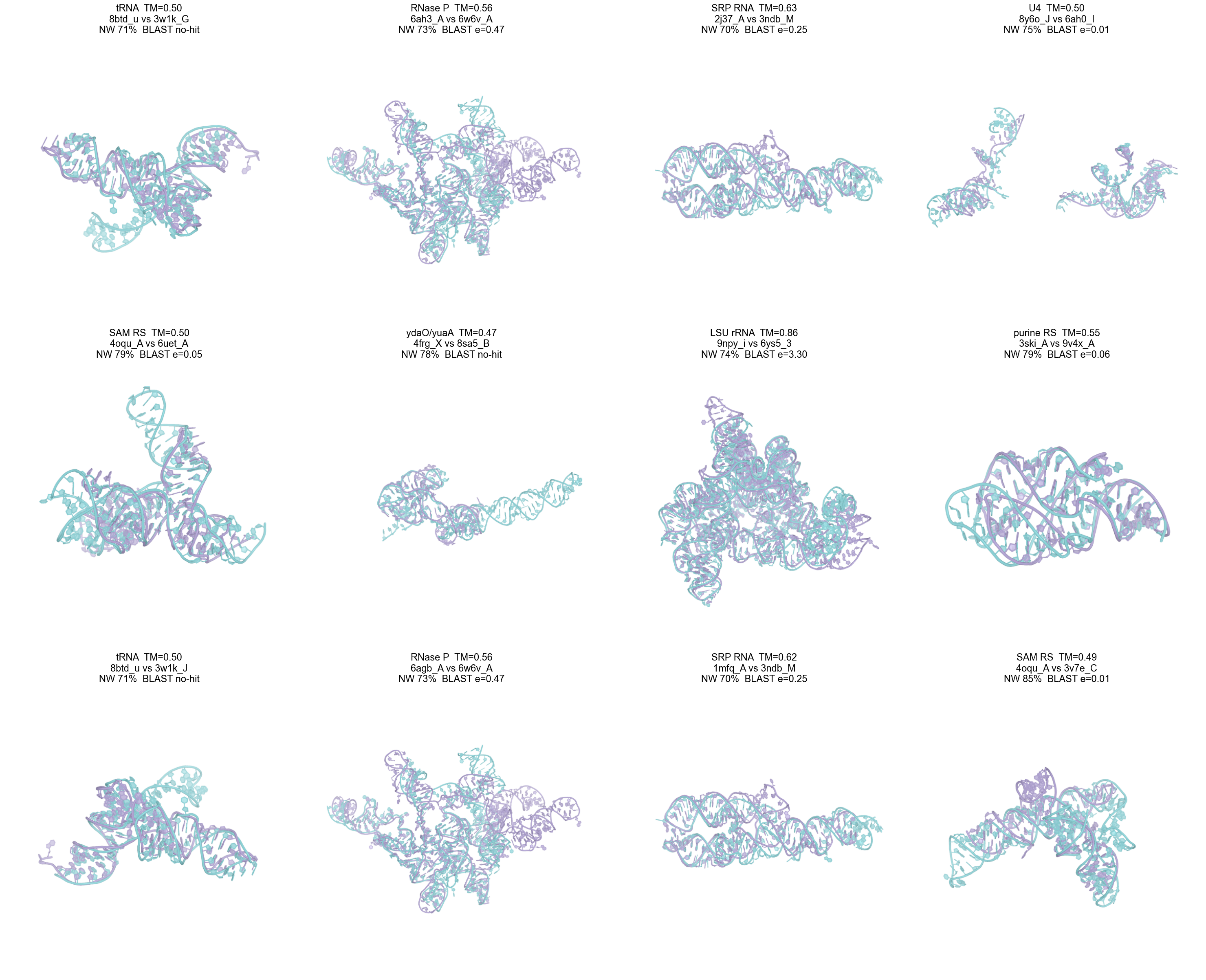

Supplementary companion to main-text Figure 4a: the same twelve cross-family within-Rfam-clan case superpositions, each panel annotated with clan name, TM-score, chain identifiers, NW pairwise identity and BLAST *e*-value.

#### Figure S9. RS-20-favoured cases view with full annotation

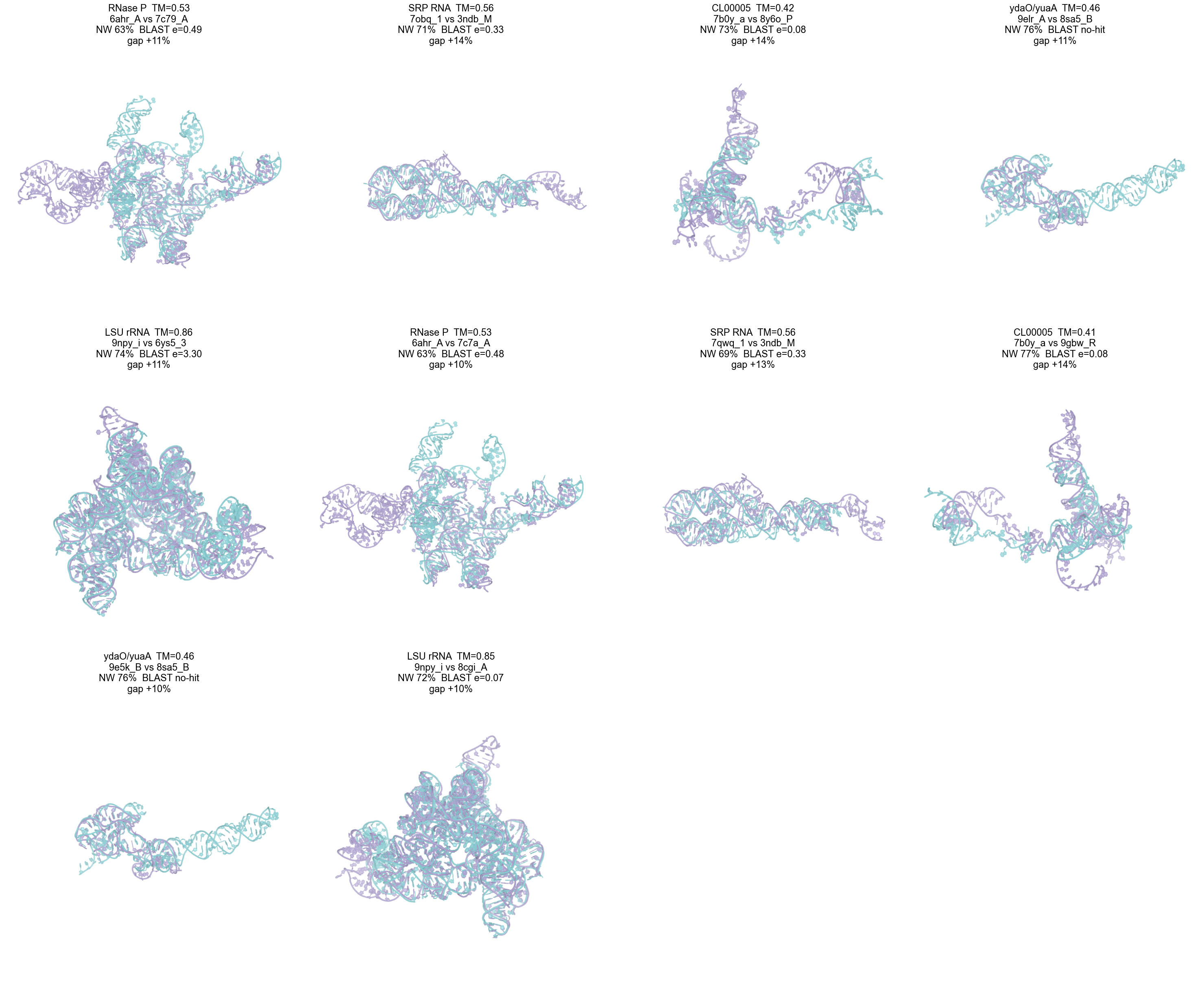

Supplementary companion to main-text Figure 4b: the same twelve representative RS-20-favoured cases in which the 20-letter RS-20 alphabet outranks the 92-letter iPARTS2 alphabet in percentile rank, now annotated with clan, TM, chain pair, NW identity, BLAST *e*-value and the percentile gap between RS-20 and iPARTS2.

#### Figure S10. Infernal complementarity

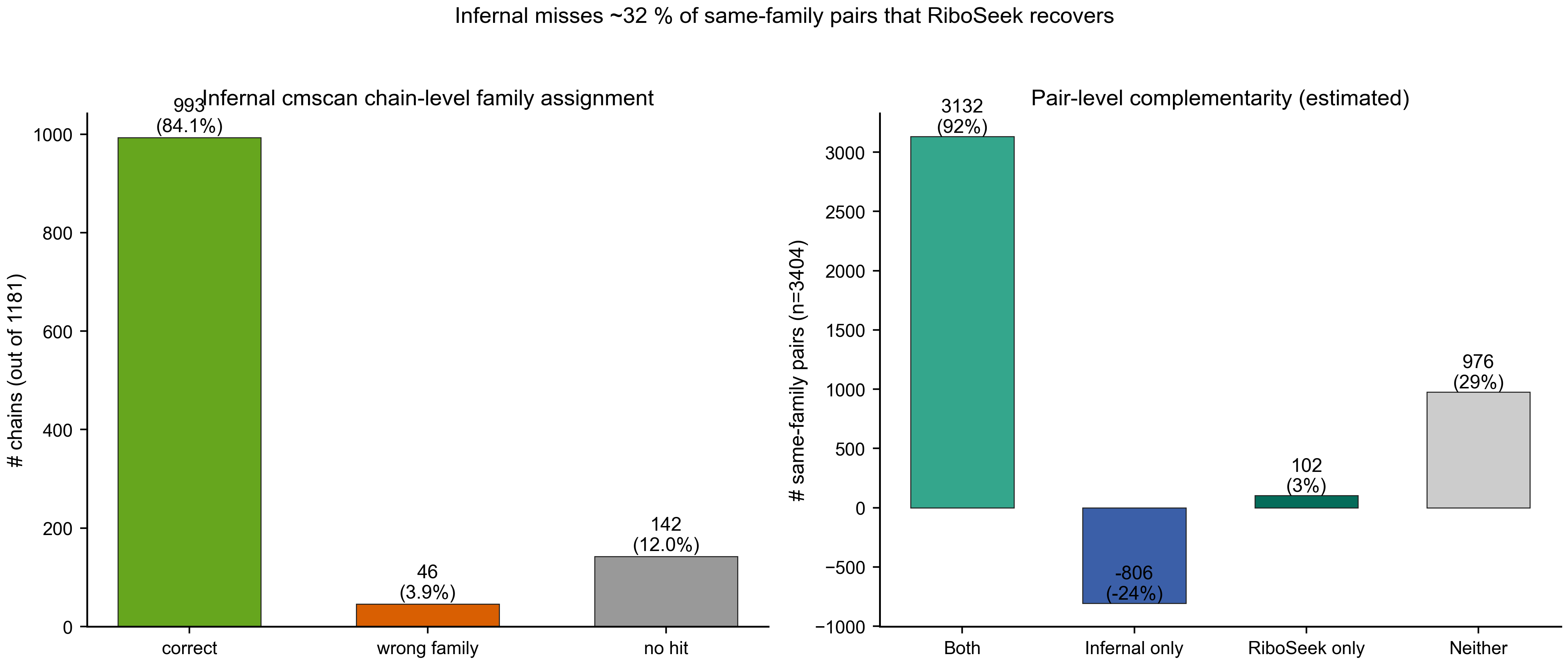

Set-overlap of same-family pairs correctly recovered by Infernal cmscan vs. RiboSeek on the benchmark. Infernal misses ~32 % of same-family pairs (chains where Infernal returns no hit, the wrong family, or where partial domains escape the covariance model); RiboSeek recovers most of these.

#### Figure S11. Combined hub-validation TM distribution (48 seeds)

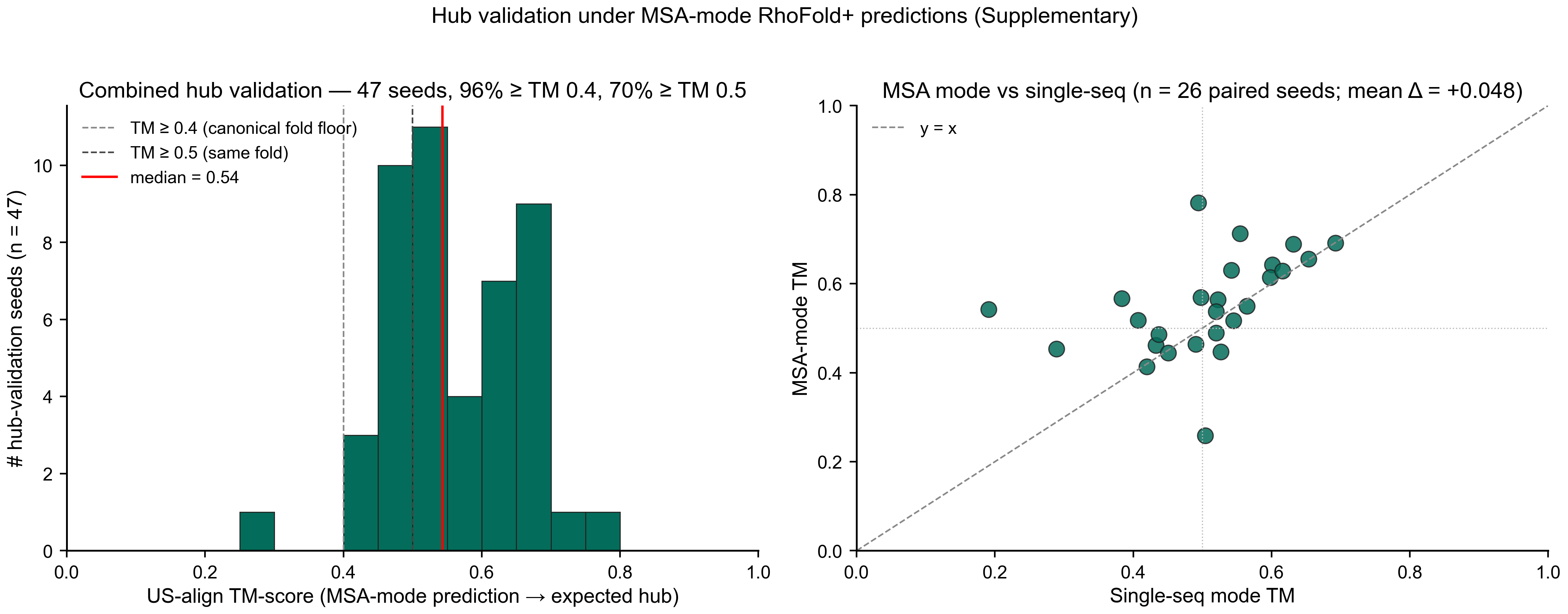

(a) Histogram of US-align TM-score for all 47 evaluable hub-validation seeds (27 original + 21 extension, all MSA-mode RhoFold+) against their expected experimental hub chain. Dashed lines at TM = 0.4 and TM = 0.5; red line = median (0.54). 98 % of seeds exceed TM 0.4; 70 % exceed TM 0.5. (b) Paired scatter: MSA-mode TM (y-axis) vs. single-sequence-mode TM (x-axis) for the 26 seeds with both measurements. Points above the diagonal indicate MSA-mode improvement; mean gain = +0.048 TM.

### Supplementary Tables

#### Table S1. Feature vectors of published RNA structural alphabets

| Alphabet | Year | Reference | Letters | Feature vector (per-residue) | Spatial neighbours encoded? |
| --- | --- | --- | --- | --- | --- |
| SARSA | 2008 | Chang et al.[5] | 23 | 4 backbone torsions (α, γ, δ, ζ) | No |
| iPARTS | 2010 | Wang et al.[6] | 23 | 2 pseudo-torsions (η, θ) | No |
| R3D-BLAST | 2011 | Liu et al.[8] | 23 | 2 pseudo-torsions (η, θ) | No |
| iPARTS2 | 2016 | Yang et al.[7] | 92 | 2 pseudo-torsions + base identity | No |
| R3D-BLAST2 | 2017 | Yen et al.[9] | 23 | 2 pseudo-torsions (η, θ) | No |
| DNATCO | 2019 | Xiong et al.[19] | ≈12 | 12 dinucleotide-level geometric parameters | No (local only) |
| **RS-20 (this work)** | 2026 | — | **20** | **top-3 spatial neighbours × 3 features + base orient. + seq dist. (14 D)** | **Yes** |
| **RS-80 (this work)** | 2026 | — | **80** | **RS-20 × nucleotide identity** | **Yes** |

#### Table S2. 29 rendered 3D superposition cases (coverage view, Fig. 4a)

The full table of chain IDs, Rfam families, clan, TM, NW %id, BLAST e-value, method percentile ranks and chain lengths for each of the 29 coverage-view cases is provided in the supplementary data file case_superpositions.json.

#### Table S3. Software and hardware

| Software | Version |
| --- | --- |
| Python | 3.11 (main env), 3.8 (RhoFold+ env) |
| scikit-learn | 1.3.0 |
| Biopython | 1.79 |
| PyTorch | 2.5.1+cu124 (main), 1.10.2+cu113 (RhoFold+) |
| US-align | 20260329 |
| NCBI BLAST+ | 2.16.0 |
| Foldseek | latest as of April 2026 |
| RhoFold+ | commit from ml4bio/RhoFold (April 2026) |
| **Hardware** | 8 × NVIDIA A100 80 GB PCIe; single-threaded C NW/SW for benchmarks; 16-core multi-process for dark-family search |

#### Table S4. Foldseek pseudo-protein control: mapping sensitivity

See Supplementary Methods § S4 for full discussion.

| Mapping | A → | U → | C → | G → | Overall AUC | no-hit rate | high-sim AUC |
| --- | --- | --- | --- | --- | --- | --- | --- |
| M1 mnemonic | Ala | Val | Cys | Gly | 0.8666 | 45.86 % | 0.179 |
| M2 size | Trp | Gly | Ala | Phe | 0.8675 | 46.05 % | 0.179 |
| M3 polarity | Arg | Thr | Ser | Lys | 0.8660 | 46.23 % | 0.170 |
| M4 random | Asp | Glu | Leu | Ile | 0.8664 | 46.13 % | 0.176 |

All four mappings are within 0.0015 AUC of each other (spread 0.8660– 0.8675), with no-hit rates within 0.4 pp (45.86–46.23 %) and high- similarity AUCs within 0.01 (0.170–0.179). The failure pattern is a property of the 3Di encoder’s geometric training distribution, not of the residue-naming choice. The M4 random-mapping control reproduces the mnemonic-mapping signature within noise, which specifically addresses “why these particular amino-acid assignments.”

#### Table S5. Dark-family pilot summary

| Quantity | Count |
| --- | --- |
| Rfam families, total (v14.10) | 4,227 |
| Rfam families with ≥ 1 PDB chain in our data | 157 |
| Rfam families without PDB representation (dark) | 4,070 |
| Dark families with 40–200 nt AUGC-only seed | 3,694 |
| RhoFold+ predictions completed | 3,680 (99.6 %) |
| Valid RS-20 encodings (≥ 10 letters) | 3,680 |
| Top-100 dark queries US-align verified | 100 |
| Cross-family dark cluster (TM ≥ 0.40, different Rfam) | 8 |
| Rfam annotation-gap hit (TM ≥ 0.40, unlabeled target) | 41 |
| Dark hubs (≥ 3 dark families → single target) | 5 |
| Hub validation seeds (additional, independent) | 27 |
| Hub validation weighted top-1 recall | 63 % |

#### Table S6. Eight cross-family dark clusters (Fig. 5e)

| Dark family (query) | Experimental target | Target Rfam | TM | RS-80 | Seq NW %id |
| --- | --- | --- | --- | --- | --- |
| RF01013 | 8fcs_A | RF00661 (mir-31) | **0.681** | 2.74 | 0.79 |
| RF02019 | 8fcs_A | RF00661 (mir-31) | 0.584 | 2.83 | 0.79 |
| RF03230 | 8fcs_A | RF00661 (mir-31) | 0.550 | 2.71 | 0.79 |
| RF03588 | 9obm_A | RF00051 (miR-17/92) | 0.522 | 2.84 | 0.74 |
| RF03224 | 2l3j_B | RF02095 | 0.517 | 2.82 | 0.71 |
| RF02944 | 8v1h_A | RF01684 | 0.445 | 2.73 | 0.78 |
| RF04236 | 9de7_A | RF00250 | 0.410 | 2.73 | 0.58 |
| RF03339 | 2kuw_A | RF00207 | 0.407 | 2.85 | 0.67 |

#### Table S7. Five dark hubs with member dark families

| Hub (experimental chain) | Target Rfam | Converging dark Rfam families | n seeds tested | Weighted top-1 recall |
| --- | --- | --- | --- | --- |
| 8scz_B | — (unannotated) | RF00588, RF01944, RF04270, RF03970, RF03819, RF03652 | 9 | 33 % |
| 9kby_B | — (unannotated) | RF04105, RF03703, RF04242, RF03857, RF00914 | 5 | **100 %** |
| 8fcs_A | RF00661 (mir-31) | RF02019, RF01013, RF03230 | 5 | 80 % |
| 7kub_A | — (unannotated) | RF03418, RF03359, RF03293, RF03479 | 5 | 80 % |
| 8dg5_E | — (unannotated) | RF03431, RF00955, RF03865 | 3 | 33 % |
| **Weighted overall** |  |  | **27** | **63 %** |

#### Table S8. Cross-family within-clan case list (387 pairs)

The complete list of 387 clan-guided cross-family within-Rfam-clan pairs with US-align TM ≥ 0.4 and BLAST *e*-value > 0.01 — comprising chain identifiers, Rfam families, clan, TM, BLAST *e*-value, NW pairwise identity, and RS-20 / RS-80 / iPARTS2 percentile ranks — is provided as machine-readable supplementary data in case_studies_clan_guided.json.

#### Table S9. Computational footprint

| Resource | Value |
| --- | --- |
| RhoFold+ dark-family predictions (3,694 families) | ~6 GPU-hours on 8 × A100 |
| K-means training (15-D, K = 20) | ~30 min (single CPU) |
| Hard-negative benchmark (11,170 pairs, 6 methods) | ~12 min |
| US-align verification (dark-family top 100) | ~30 min |
| Alphabet + score matrix files | ~5 MB |
| 3,694 predicted PDBs | ~500 MB |
| Enhanced features (15,391 chains) | ~1.3 GB |

#### Table S10. Feature-ablation AUC values

Bootstrap 95 % confidence intervals (n = 1,000 resamples) on NW+SW AUC for nested feature configurations, under the family-held-out K-means protocol described in Methods § Family- aware K-means. Referenced in main-text Fig. 2a.

| Configuration | n features | AUC [95 % CI] |
| --- | --- | --- |
| Backbone only (η, θ) | 2 | 0.901 [0.894, 0.907] |
| Backbone + base orientation | 4 | 0.907 [0.901, 0.913] |
| Backbone + sequential distances | 5 | 0.916 [0.910, 0.922] |
| All 7 traditional features (no neighbour) | 7 | 0.921 [0.915, 0.926] |
| **3 spatial-neighbour features alone** | **3** | **0.945 [0.940, 0.950]** |
| All 10 features (traditional + neighbour) | 10 | 0.946 [0.942, 0.951] |
| 15-D drop-backbone (RS-20 production) | 15 | 0.955 [0.951, 0.959] |

### Supplementary Text

#### The “BLAST twilight zone” and why sequence identity breaks down on RNA

BLAST’s seed-and-extend heuristic detects homology by finding a short exact or near-exact match (typically 7 nucleotides for blastn default), extending that seed greedily to find an aligned region of sufficient score. This works well when:

(i) the alignment contains long contiguous high-identity stretches (the seed can be placed with high probability); and (ii) the alphabet is large enough that random 7-mer matches are rare (4⁷ = 16 K possible sequences).

Both conditions fail for cross-family RNA homologs at 50–70 % pairwise identity:

- On a 100-nt sequence at 60 % identity, the probability of any single 7-mer being perfectly conserved is 0.6⁷ = 2.8 %; even on a 100-nt pair, the expected number of such seeds is only 2.8, and they are often broken up by intervening mismatches.
- RNA has a 4-letter alphabet (vs. protein’s 20), so random chance matches are already 16× more frequent per length unit; background noise dominates signal in precisely the regime where structural homology persists.
- Short functional RNAs (30–500 nt) have few absolute seeds to work with.

Empirically (Fig. 3b, main), BLAST AUC on our benchmark is 0.60 in the NW 50–70 % stratum — virtually random. RS-80 stays at 0.95 because structural conservation is still strong at 50 % sequence divergence and the alphabet encoding is immune to the seed-and- extend heuristic’s failure.

#### Why the Rfam family structure is finer than the structural super-family

Rfam families are defined by covariance models (CMs) — profile models that capture position-specific base preferences and co-variation signal indicative of conserved secondary structure. Two RNAs belong to the same Rfam family if both score above the family’s CM cutoff threshold. This definition captures:

(i) sequence conservation at conserved sites (e.g., active-site bases in ribozymes); (ii) co-variation patterns indicative of base-pairing (indirect 2D structure conservation); (iii) conserved length and overall secondary structure topology.

What CMs **do not directly capture** is 3D structural super-family identity: two families with different mature-sequence cores but similar global fold (e.g., multiple microRNA precursor Rfam families all forming a similar hairpin shape with different mature miRNA sequences) will not cross-hit on each other’s CMs. Our dark-hub observation (Fig. 5d) reflects this directly: 5 PDB chains attract ≥ 3 dark Rfam families, indicating structural super-family membership that Rfam’s sequence-based CM partition does not express.

This is not a criticism of Rfam, which does not claim to capture structural super-family identity; Rfam’s mandate is sequence family annotation. Rather, it identifies a specific use case for a structure-based search engine: predicting structural super-family membership where it cuts across Rfam’s sequence-level partition.
